# PlasmoFP: leveraging deep learning to predict protein function of uncharacterized proteins across the malaria parasite genus

**DOI:** 10.1101/2025.09.12.675843

**Authors:** Harsh R. Srivastava, Daniel Berenberg, Omar Qassab, Ziyi Wang, Richard Bonneau, Jane M. Carlton

## Abstract

The first malaria parasite *Plasmodium falciparum* genome published in 2002 jump-started functional studies, but a large fraction of all predicted proteins remains partially annotated and of ‘unknown function’. Here, we introduce Plasmodium Function Predictor (PlasmoFP), deep learning models designed specifically for species of genus *Plasmodium*. Innovatively, PlasmoFP models are trained on structure-function relationships of the phylogenetically relevant SAR (Stramenopiles, Alveolate, and Rhizarians) supergroup proteins, addressing challenges to annotating *Plasmodium* proteins due to their low sequence similarity to well-characterized model organism proteins. PlasmoFP models estimate epistemic uncertainty, control false discovery rates in model predictions, and are validated using proteins with manually curated GO terms and experimentally characterized proteins. Integrating PlasmoFP predictions with current protein annotations, we reduced the proportion of unannotated proteins without Gene Ontology terms from 15-59% to 3-28% across 19 *Plasmodium* species, and improved the proportion of fully annotated proteins from 7-42% to 36-68%. PlasmoFP predictions advance *Plasmodium* basic research, an important component of global malaria R&D.

## Introduction

Malaria remains a significant global health burden, with an estimated 263 million cases reported in 2023 across 83 endemic countries (*1*). Infections are caused by protozoan parasites of genus *Plasmodium*, a member of phylum Apicomplexa within clade Alveolata, which together with Stramenopiles and Rhizaria forms the Stramenopiles–Alveolates-Rhizaria (SAR) eukaryotic supergroup (*2*). Five major species are known to cause most human malaria: *P. falciparum*, *P. vivax*, *P. malariae, P. ovale,* and *P. knowlesi* (*1*). Other *Plasmodium* species that are used as human disease models infect mammals include African tree-dwelling rodents (e.g., *P. berghei*) and non-human primates (e.g., *P. coatneyi*), while more evolutionarily distant species infect birds and lizards (e.g., *P. gallinaceum*) (*3*). Since the publication of the first *Plasmodium* genomes in 2002 (*P. falciparum*(*4*) and *P. yoelii* (*5*)), successive efforts have generated assemblies of species from virtually every major clade within the genus. This information is consolidated in PlasmoDB, a resource that provides standardized genomic, transcriptomic, and functional annotations across *Plasmodium* species (*6*).

*Plasmodium* genome sequencing has outpaced gene functional annotation, with many predicted gene products labeled simply as “proteins of unknown function” (PUFs). Even in the extensively studied *P. falciparum* proteome, 16% of protein coding genes lack any annotation from the tripartite Gene Ontology (GO) vocabulary of molecular function (MF), biological process (BP), and cellular component (CC), while 42% are partially GO annotate. In the less well characterized *P. ovale curtisi*, ∼50% of proteins are unannotated, 38% are partially annotated, and just 12% possess full GO annotations (*7*). Inadequately characterized *Plasmodium* proteins constitute a major bottleneck to malaria parasite basic research, impeding efforts to elucidate parasite survival strategies and host–parasite interactions.

The functions of ‘unknown’ proteins have traditionally been elucidated through resource-intensive wet-lab experiments(*8*), which are an impractical method for high-throughput proteome-wide studies in challenging non-model organisms like *Plasmodium*(*9*). Automated function prediction (AFP) techniques offer an alternative by leveraging the relationship between protein sequence, structure, and function to infer annotations. AFP annotations often encompass family- or ontology-based classifications such as Pfam domains, InterPro signatures, and GO terms(*10-12*); the latter are widely used in multi-omics analyses to uncover mechanistic processes underlying experimental results(*11, 13*).

Many AFP methods use sequence similarity approaches such as BLAST, PSI-BLAST, and HMMER, which infer protein function by comparing query sequences to protein family models associated with characterized functions, or well-curated proteins in databases like SwissProt(*10, 14-17*). In parallel, early AFP approaches employed genetic interaction networks derived from large scale genomic interrogation of model species and well-studied organisms chosen for their low cost and tractable genetics (*18, 19*). Not surprisingly, these methods are relatively ineffective for proteins with low sequence similarity to known proteins(*20-22*). A significant proportion of *Plasmodium* proteomes is either unique to the genus, or is shared only with other poorly-annotated apicomplexan parasites, or is only remotely homologous to well-characterized proteins(*4*). Additionally, *Plasmodium* proteins frequently contain long intrinsically disordered regions which do not align well, further reducing the performance of sequence-based approaches(*23*). Such limitations explain why many *Plasmodium* proteins remain functionally uncharacterized and underscore the need for alternative strategies.

Recent advances in deep learning have transformed AFP by enabling models to extract signal directly from protein sequences and capture heuristics correlated with protein sequence, structure, and function(*17, 24-28*). Foundational protein language models, such as ProtT5 and ESM, have shown great promise in encoding sequence- and structure-based features into information-rich, relatively low dimensional vector embeddings(*29, 30*). These advances provide groundwork for exploring alternative strategies beyond traditional sequence similarity approaches. We hypothesize that leveraging the direct relationship between protein structure and function while jointly conditioning learning on phylogenetically relevant relationships offers a more effective strategy for predicting the functions of highly divergent proteins like those found in *Plasmodium*.

Here, we introduce *Plasmodium* Function Predictor (PlasmoFP), a family of deep learning-based AFP models trained separately on each of the three GO subontologies to predict GO terms for *Plasmodium* PUFs and partially annotated proteins. PlasmoFP explicitly models phylogenetically relevant relationships between structure and function within the SAR taxonomic supergroup. Through comprehensive evaluation of model performance and ablations of model architectures, we demonstrate that PlasmoFP outperforms both baseline AFP approaches and existing AFP methods on *Plasmodium* protein annotations that have been manually curated. We additionally show PlasmoFP predicts meaningful GO terms for a set of experimentally characterized RNA-associated proteins in *P. falciparum,* as well as sustains performance on proteins with high levels of intrinsic disorder. In turn, PlasmoFP represents the most effective function prediction model developed specifically for the genus *Plasmodium*. We show that integrating PlasmoFP predictions with existing annotations across the MF, BP, and CC GO subontologies significantly reduces the proportion of *Plasmodium* PUFs and increases the proportion of proteins annotated with all three GO subontologies. We also demonstrate that PlasmoFP predictions refine and expand important RNA-associated proteins in the genus, and uncover new Plasmodium biology through expanding protease-like and transport/localization protein repertoires in malaria parasites that primarily infect humans. Collectively, PlasmoFP predictions offer a critical resource for advancing our understanding of *Plasmodium* biology and serve as a foundation for the basic research required for development of novel therapeutic and diagnostic strategies for malaria.

## Results

### PlasmoFP models are trained to predict GO terms for each subontology

We developed PlasmoFP, three independently trained deep-learning models on protein structure–function relationships, one for each GO subontology (MF, BP, and CC). Our PlasmoFP models use TM-Vec embeddings(*31*), structure-imbued vectors of fixed dimensionality, to predict protein function. TM-Vec fine-tunes an underlying sequence encoder (e.g., ProtT5) with a structural similarity learning objective and has been shown to perform well in several tasks related to AFP(*29, 31*). Motivated by the evident expressive power and versatility of TM-Vec embeddings, we followed previous work in AFP by choosing to model protein function by *Pθ(y | xs)*, where *θ* is the set of learnable parameters of a multi-layer perceptron (MLP), *y* ∈ {0, 1}^c^ is a vector of GO term annotations, and *xs* ∈ R^d^ is the TM-Vec embedding of a protein sequence *s* **(Fig. 1)**. PlasmoFP models are trained to optimize the binary cross-entropy objective, with no gradient updates applied to the TM-Vec encoder. Our choice to keep the TM-Vec model frozen during training forces PlasmoFP to learn structure-function relationships from a fixed-sequence structure prior with a relatively small number of parameters rather than requiring large-scale updates to the underlying protein language model. PlasmoFP models are trained on automatic and manual assertion GO terms (see Methods), with sequences partitioned by similarity within the SAR taxonomic supergroup to prevent information leakage. We evaluated models on a SAR test set and a curated *Plasmodium* holdout set consisting of manually curated GO annotations held out entirely from model training **(Fig. S1)**.

**Fig. 1.**
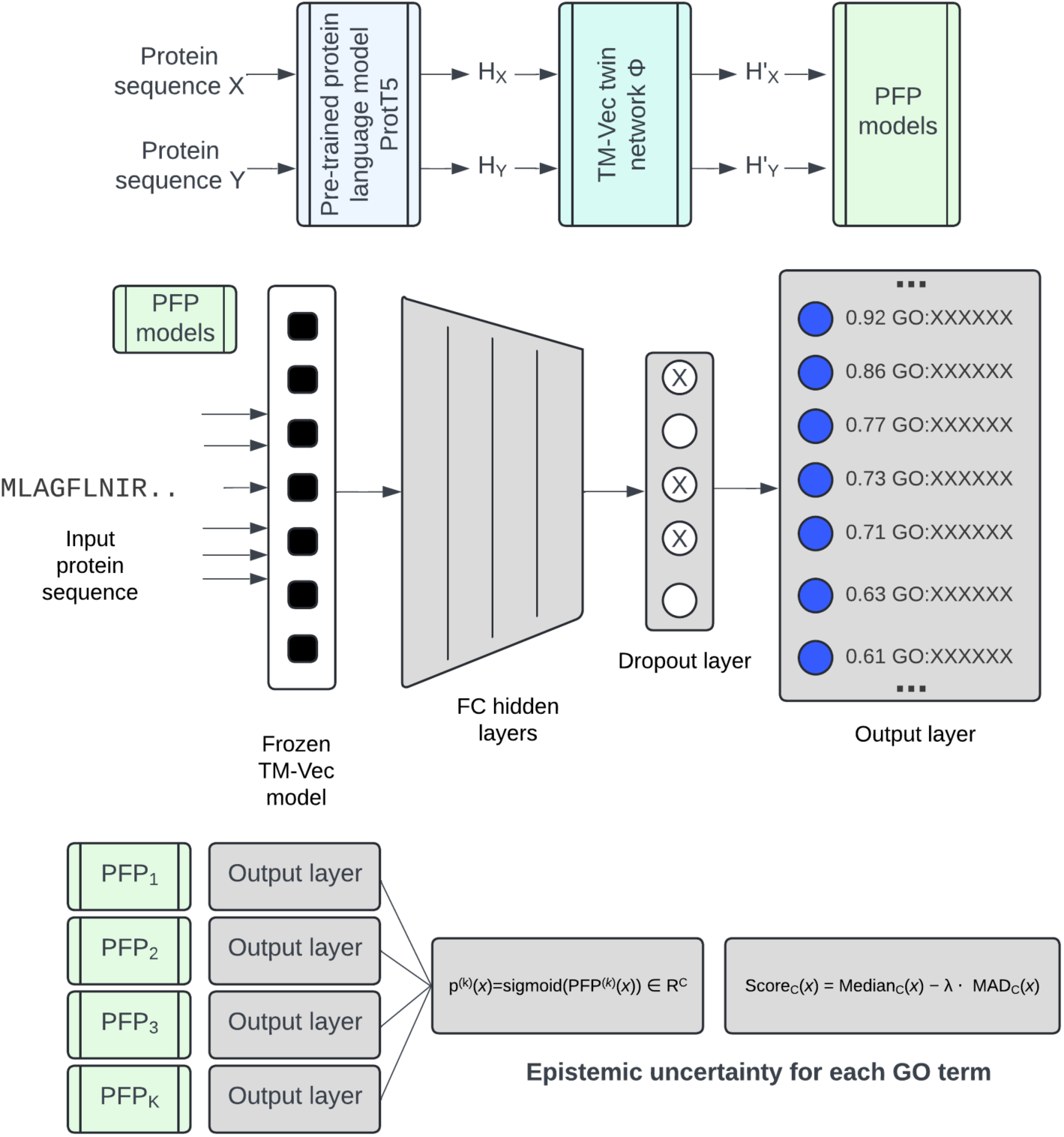
PlasmoFP models train on structure representations and utilize deep ensembles. **(A)** Schematic representation of the workflow for training PlasmoFP models using protein language model embeddings. Protein sequences (e.g., X and Y) are processed through the pre-trained ProtT5 language model to generate embeddings (HX and HY). These embeddings are then fine-tuned through the TM-Vec twin network (Φ) to produce sequence-specific representations (H’X and H’Y), which serve as inputs to the PlasmoFP models during model training and inference. **(B)** PlasmoFP models predict GO term probabilities from the processed sequence embeddings (H′) via fully connected (FC) layers. (C) Each PlasmoFP model for each subontology is configured as a deep ensemble trained on different *k* folds of the training dataset. We calculate per-term epistemic uncertainty and create a conservative uncertainty adjusted probability for each input-GO term pair.

### PlasmoFP models quantify uncertainty with deep ensembles

For each of the three GO subontologies (MF, BP, and CC), we used four PlasmoPF models to capture epistemic uncertainty, the reducible component of error due to limited model knowledge(*32*). We evaluated four approximations to the posterior predictive, *i.e.*, the distribution of outcomes after averaging over many plausible models learned from the data: (1) Monte Carlo Dropout (MCD)(*33*); (2) a *k-*member temporal ensemble (predictions from the last *k* checkpoints); (3) a *k*-fold ensemble (*k* independently initialized models); and (4) a deterministic baseline with no uncertainty **(Fig. S2, Fig. 1B)**. For each method, we then calculated uncertainty-adjusted model probabilities, *i.e.*, central estimate minus dispersion, to curb overconfident errors **(Fig. S3)**.

Across all subontologies, the performance of the temporal ensemble behaved nearly identically to the deterministic baseline **(Fig. 2A)**. This indicates that successive checkpoints are highly correlated and therefore fail to provide meaningful epistemic uncertainty across *k* checkpoints. In contrast, increasing the number of MCD passes led to a gradual drop in (false discovery rate) FDRmicro and sharpness, and an increase in PrecisionMacro **(Fig. 2A)**. This suggests additional stochastic samples help the model approximate the posterior predictive distribution more faithfully while producing less overconfident predictions. Deep ensembles exhibited a similar trend; as *k* increased, FDRmicro decreased and PrecisionMacro increased **(Fig. 2A)**. Notably, at *k* = 20, we observed the largest gain in PrecisionMacro without substantial decrease in RecallMacro. This implies that independent weight initializations explore diverse local optima, providing a richer posterior approximation that yields fewer false positives. Given the results of this analysis, we selected *k* = 20 deep ensembles for all downstream inference.

**Fig. 2.**
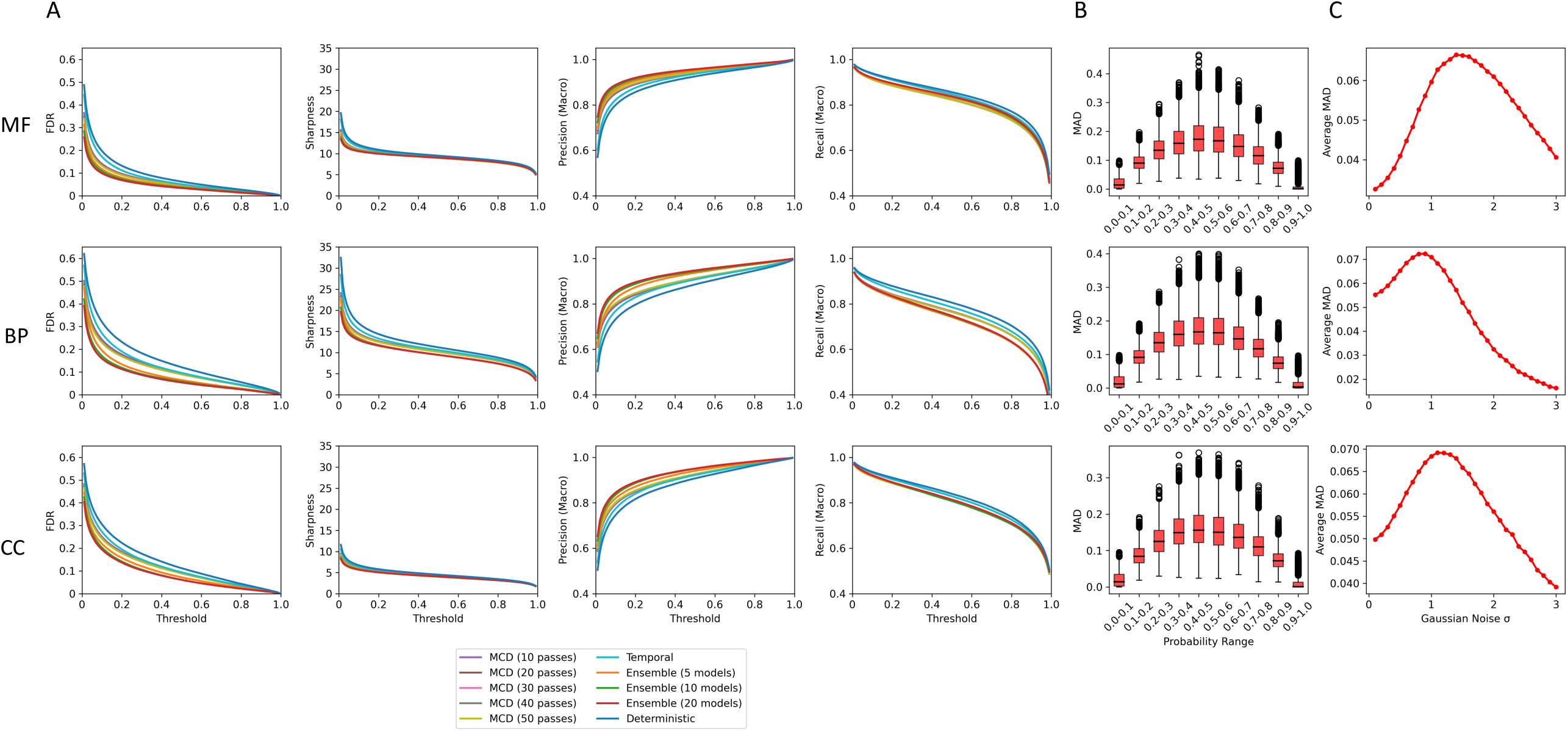
Deep ensembles improve model precision, FDR and sharpness. **(A)** FDR (micro), sharpness, precision (macro) and recall (macro) curves for across all the subontologies for all models. **(B)** Boxplots showing the distribution of MAD for true positive predictions at binned softmax probabilities. **(C)** Average MAD of true positive predictions at differing levels of Gaussian noise added to input TM-Vec embeddings.

We next examined the correlation between uncertainty and each PlasmoFP model’s softmax confidence and found that predictions with softmax probability around 0.5 exhibited the highest uncertainty **(Fig. 2B)**. We also observed an inverse correlation between the log count of true positives per GO term in the test set and epistemic uncertainty, not only suggesting that rarer terms with fewer positive examples are ascribed higher uncertainty, but also indicating that our estimates capture data scarcity and underrepresentation during model training **(Fig. S4)**. For PlasmoFP GO MF predictions, the highest uncertainty was observed for enzyme-related functions, including lyases, ligases, and transferases **(Fig. S5)**. These functions also tended to have low to medium information accretion values, suggesting that the model struggles most with functions requiring fine-grained, residue-level features that are not fully captured by per-protein embeddings. To further validate our uncertainty estimates, we perturbed TM-Vec embeddings with Gaussian noise and found that uncertainty initially rose as inputs drifted from the training manifold, before declining at higher noise levels when ensemble members converged on uniformly low-confidence predictions **(Fig. 2C)**.

Most existing function-prediction methods select a single decision threshold to maximize Fmax on a given dataset. We introduce an alternative approach that uses empirically observed per-term FDR on the SAR test set to assemble predicted GO term sets at specified FDR thresholds for each PlasmoFP model. We adopt the conservative term “expected FDR” (eFDR) at inference time. *Plasmodium* PUFs are included in the SAR supergroup, so these empirically-derived eFDR thresholds should generalize to *Plasmodium* proteins. This method directly controls the expected FDR of prediction sets by tailoring uncertainty-adjusted threshold cutoffs to each term rather than balancing overall recall and precision with a one-size-fits-all threshold. We find that the per-term variability in uncertainty-adjusted probabilities required to meet an FDR cutoff is substantial **(Fig. S6)**. At stringent FDR values (e.g., 1–5% eFDR), some GO terms require uncertainty-adjusted probability cutoffs above 0.9, while others can be called at around 0.5. Notably, CC terms are the most conservative, demanding higher uncertainty-adjusted probabilities, whereas MF terms can be predicted at much lower uncertainty-adjusted probabilities for the same error rate **(Fig. S6).**

By performing a grid search over eFDR levels **(Fig. S7)**, we identified an optimal range of 10-20% eFDR within which Recallmicro remains high without large decreases in Precisionmicro. Beyond 30% eFDR, recall gains plateau at the cost of precision. Smin, an information-content weighted metric combining *remaining uncertainty* (ground truth terms not predicted) *(ru)* and *misinformation* (false positive predictions) *(mi),* reveals a convex trade-off as *ru* falls rapidly as cutoffs relax, while *mi* climbs linearly. This yields an Smin minimum also at 10–20% eFDR across all subontologies.

### Phylogenetic relevance and TM-Vec embeddings improve function prediction in SAR

We evaluated the impact of conditioning model training on phylogenetically related sequences and incorporating structural information, on function-prediction performance within the SAR supergroup. For each GO subontology, we compared baseline methods along two orthogonal axes. First, to isolate the data effect, we held the algorithm constant while varying the training set, comparing (1) CAFA-CNN, trained on CAFA-5 SwissProt sequences, versus SAR-CNN, trained on SAR-specific sequences; (2) CAFA-TM-Vec trained on TM-Vec embeddings for CAFA-5 SwissProt sequences set versus PlasmoFP, trained on TM-Vec embeddings for SAR-specific sequences; (3) BLAST against SwissProt versus BLAST against SAR-specific sequences; (4) Foldseek against SwissProt versus Foldseek against SAR-specific sequences(*14, 34*) (**Fig. S8)**. In all comparisons, identical train and test splits were used to ensure that observed differences in Fmax and Smin reflect purely the impact of phylogenetic conditioning during training or the use of structure-imbued embeddings.

The inclusion of phylogenetically relevant sequences during training improved performance across architectures: SAR-CNN achieved higher Fmax and lower Smin than CAFA-CNN; PlasmoFP models outperformed CAFA-TM-Vec models; and BLASTTrain outperformed BLASTSwissProt in all subontologies **(Table 1)**. We observed no significant hits when querying SAR test sets with FoldseekSwissProt, underscoring the scarcity of detectable structure-based homologs to the broader SwissProt set. Taken together, these results indicate that restricting training or search data to phylogenetically relevant sequences improves model accuracy.

**Table 1.** PlasmoFP performance on SAR test set. Bolded values indicate best performance in each subontology; FMacro values are FMacro | max for all models except PlasmoFP 10% eFDR. Smin values are reported at the threshold corresponding to FMacro | max.

| | Model | $F_{\text{Macro}}$ | $S_{\text{min}}$ |
| --- | --- | --- | --- |
| MF | **PlasmoFP 10% eFDR** | **0.900** | **2.050** |
|  | CAFA-TM-Vec | 0.632 | 7.425 |
|  | SAR-CNN | 0.721 | 5.514 |
|  | CAFA-CNN | 0.456 | 10.062 |
|  | BLAST<sub>pSwissProt</sub> | 0.639 | 24.525 |
|  | BLAST<sub>pTrain</sub> | 0.765 | 9.328 |
|  | Foldseek<sub>Train</sub> | 0.844 | 10.613 |
| BP | **PlasmoFP 10% eFDR** | **0.851** | **3.780** |
|  | CAFA-TM-Vec | 0.593 | 8.826 |
|  | SAR-CNN | 0.644 | 7.466 |
|  | CAFA-CNN | 0.396 | 12.415 |
|  | BLAST<sub>pSwissProt</sub> | 0.514 | 57.571 |
|  | BLAST<sub>pTrain</sub> | 0.782 | 9.827 |
|  | Foldseek<sub>Train</sub> | 0.764 | 12.329 |
| CC | **PlasmoFP 10% eFDR** | **0.875** | **1.235** |
|  | CAFA-TM-Vec | 0.698 | 2.864 |
|  | SAR-CNN | 0.766 | 2.175 |
|  | CAFA-CNN | 0.687 | 2.918 |
|  | BLAST<sub>pSwissProt</sub> | 0.501 | 20.343 |
|  | BLAST<sub>pTrain</sub> | 0.777 | 3.008 |
|  | Foldseek<sub>Train</sub> | 0.771 | 3.900 |

Using a fixed training set to isolate the model effect, we varied the method comparing (1) CAFA-CNN versus CAFA-TM-Vec; (2) SAR-CNN versus PlasmoFP; and (3) BLAST versus Foldseek on SAR databases. We found that training or searching using structure-imbued representations enhanced performance, as reflected by higher Fmax and lower Smin when comparing CAFA-TM-Vec to CAFA-CNN, and PlasmoFP to SAR-CNN across all subontologies **(Table 1)**. When comparing FoldseekTrain to BLASTTrain we observed higher F-scoreMacro results for the MF GO ontology and comparable performance for the BP and CC ontologies, consistent with MF annotations being more tightly linked to local structure-based motifs than the broader, protein-level biology captured by BP and CC. Taken together, these results indicate that explicitly leveraging structure–function relationships helps address remote homology in sequence space, consistent with prior work showing structure is highly informative for functional characterization in *P. falciparum*(*35*). Additionally, the larger performance gain of PlasmoFP models over FoldseekTrain implies that structure-imbued protein embeddings can offset the impact of lower-quality predicted structures during structure–structure searches for function prediction.

To assess whether PlasmoFP models learned SAR-specific structure–function relationships, we evaluated performance on CAFA-5 proteins filtered for terms present in the PlasmoFP training data. We hypothesized that, if PlasmoFP models had specialized on SAR features, performance would decline on the broader CAFA-5 set. Consistent with this, PlasmoFP MF showed a decrease in Fmicro from 0.908 to 0.485 at 10% eFDR and an increase in Smin by ∼7 points on the CAFA-5 MF subset compared to the SAR test set **(Table 2)**. This drop was even more pronounced for BP, where Fmicro fell from 0.831 to 0.332 and Smin rose by ∼20 points at 20% eFDR, indicating that BP annotations are the most SAR-restricted. By contrast, CC was the most transferable: Fmicro declined from 0.852 to 0.537 and Smin by ∼5 points on CAFA-5. Together, these patterns suggest that PlasmoFP captures SAR-specific structure–function signals, most tightly for BP, moderately for MF, and least for CC, reflecting the deeper conservation of enzymes and core metabolism functions in the MF subontology and cellular component annotations across phylogenetic groups.

**Table 2.** PlasmoFP performance on SAR test set and CAFA test set. Bolded values indicate best performance in each subontology, and optimal eFDR thresholds for each PlasmoFP model

| | Test set (eFDR) | $F_{\text{micro}}$ | $S_{\text{min}}$ | $\text{Precision}_{\text{micro}}$ | $\text{Recall}_{\text{micro}}$ | $ru$ | $mi$ |
| --- | --- | --- | --- | --- | --- | --- | --- |
| MF | PlasmoFP 5% | 0.903 | 2.629 | 0.958 | 0.855 | 2.601 | 0.379 |
|  | **PlasmoFP 10%** | **0.908** | **2.050** | **0.918** | **0.899** | **1.855** | **0.873** |
|  | PlasmoFP 20% | 0.894 | 2.208 | 0.858 | 0.933 | 1.308 | 1.779 |
|  | PlasmoFP 30% | 0.878 | 2.838 | 0.817 | 0.949 | 1.037 | 2.642 |
|  | CAFA-5 5% | 0.478 | 9.106 | 0.509 | 0.451 | 8.69 | 2.722 |
|  | CAFA-5 10% | 0.485 | 9.006 | 0.468 | 0.503 | 8.145 | 3.842 |
|  | CAFA-5 20% | 0.478 | 9.322 | 0.422 | 0.551 | 7.582 | 5.423 |
|  | CAFA-5 30% | 0.472 | 9.867 | 0.398 | 0.579 | 7.233 | 6.711 |
| BP | PlasmoFP 5% | 0.777 | 4.716 | 0.959 | 0.653 | 4.708 | 0.273 |
|  | PlasmoFP 10% | 0.821 | 3.780 | 0.92 | 0.742 | 3.721 | 0.668 |
|  | **PlasmoFP 20%** | **0.831** | **3.140** | **0.844** | **0.818** | **2.66** | **1.669** |
|  | PlasmoFP 30% | 0.814 | 3.587 | 0.778 | 0.854 | 2.173 | 2.854 |
|  | CAFA-5 5% | 0.278 | 24.874 | 0.587 | 0.182 | 24.8 | 1.908 |
|  | CAFA-5 10% | 0.311 | 24.368 | 0.526 | 0.22 | 24.195 | 2.896 |
|  | CAFA-5 20% | 0.332 | 23.941 | 0.458 | 0.26 | 23.495 | 4.6 |
|  | CAFA-5 30% | 0.340 | 23.827 | 0.415 | 0.288 | 22.981 | 6.293 |
| CC | PlasmoFP 5% | 0.780 | 1.623 | 0.966 | 0.654 | 1.622 | 0.071 |
|  | PlasmoFP 10% | 0.830 | 1.235 | 0.926 | 0.753 | 1.22 | 0.195 |
|  | **PlasmoFP 20%** | **0.852** | **0.984** | **0.859** | **0.845** | **0.839** | **0.514** |
|  | PlasmoFP 30% | 0.839 | 1.121 | 0.792 | 0.892 | 0.647 | 0.915 |
|  | CAFA-5 5% | 0.454 | 6.462 | 0.88 | 0.306 | 6.45 | 0.399 |
|  | CAFA-5 10% | 0.492 | 6.305 | 0.831 | 0.35 | 6.269 | 0.668 |
|  | CAFA-5 20% | 0.537 | 6.118 | 0.759 | 0.415 | 6.005 | 1.169 |
|  | CAFA-5 30% | 0.564 | 6.022 | 0.704 | 0.47 | 5.782 | 1.684 |

### PlasmoFP models outperform existing protein function prediction models for *Plasmodium*

On the *Plasmodium* holdout set (a set of proteins with manually curated GO terms), PlasmoFP exhibits a consistent increase in Fmicro and a decrease in Smin as the eFDR threshold is relaxed from 5% to 30% **(Table 3)**. The magnitude of Smin reduction is greater than the relative gain in Fmicro, indicating that improved coverage is accompanied by a disproportionate drop in semantic distance to the true term. This reduction is largely driven by decreasing *ru* rather than *mi*, reflecting the intended design of PlasmoFP to recover additional true annotations while minimizing false positives at higher eFDR thresholds. When compared against DeepGOPlus and ProteInfer(*36, 37*), at the Fmax threshold, PlasmoFP outperforms both baselines on Fmicro and Smin across all subontologies, including under the strictest 5% eFDR **(Table 3)**.

**Table 3.** PlasmoFP performance on *Plasmodium* holdout set. Bolded values indicate best performance in each subontology, and optimal eFDR thresholds for each PlasmoFP model.

| | Model | $F_{\text{micro}}$ | $S_{\text{min}}$ | $\text{Precision}_{\text{micro}}$ | $\text{Recall}_{\text{micro}}$ | $ru$ | $mi$ |
| --- | --- | --- | --- | --- | --- | --- | --- |
| MF | PlasmoFP 5% eFDR | 0.885 | 3.822 | 0.982 | 0.805 | 3.816 | 0.213 |
|  | PlasmoFP 10% eFDR | 0.900 | 3.067 | 0.957 | 0.850 | 3.022 | 0.518 |
|  | **PlasmoFP 20% eFDR** | **0.902** | **2.676** | **0.920** | **0.884** | **2.464** | **1.044** |
|  | PlasmoFP 30% eFDR | 0.899 | 2.567 | 0.892 | 0.906 | 2.007 | 1.600 |
|  | ProteInfer | 0.639 | 8.363 | 0.746 | 0.559 | 7.705 | 3.252 |
|  | DeepGOPlus | 0.570 | 12.240 | 0.548 | 0.594 | 3.691 | 11.671 |
| BP | PlasmoFP 5% eFDR | 0.736 | 8.157 | 0.954 | 0.599 | 8.146 | 0.417 |
|  | PlasmoFP 10% eFDR | 0.783 | 7.179 | 0.931 | 0.675 | 7.146 | 0.697 |
|  | **PlasmoFP 20% eFDR** | **0.804** | **6.258** | **0.884** | **0.738** | **6.119** | **1.312** |
|  | PlasmoFP 30% eFDR | 0.804 | 5.899 | 0.841 | 0.770 | 5.537 | 2.034 |
|  | ProteInfer | 0.439 | 12.295 | 0.598 | 0.346 | 10.606 | 6.220 |
|  | DeepGOPlus | 0.495 | 14.831 | 0.453 | 0.545 | 6.760 | 13.201 |
| CC | PlasmoFP 5% eFDR | 0.668 | 3.623 | 0.921 | 0.524 | 3.619 | 0.180 |
|  | PlasmoFP 10% eFDR | 0.717 | 3.168 | 0.877 | 0.606 | 3.149 | 0.341 |
|  | PlasmoFP 20% eFDR | 0.754 | 2.758 | 0.838 | 0.685 | 2.702 | 0.551 |
|  | **PlasmoFP 30% eFDR** | **0.774** | **2.493** | **0.810** | **0.741** | **2.364** | **0.793** |
|  | ProteInfer | 0.437 | 4.456 | 0.599 | 0.344 | 4.026 | 1.908 |
|  | DeepGOPlus | 0.420 | 9.561 | 0.327 | 0.589 | 2.757 | 9.155 |

### PlasmoFP sustains performance with intrinsically disordered proteins

Since intrinsic disorder is widespread in *Plasmodium* proteins, we stratified SAR test sets into low (0.0–0.3), medium (0.3–0.6) and high (≥0.6) predicted disorder(*38*) and evaluated PlasmoFP at 10% eFDR **(Table 4)**. We found that overall, performance is stable: FMacro declines by 0.04 in MF and 0.05 in BP, while CC decreases by 0.03 at high disorder. Smin rises modestly by 0.37 in MF and 0.59 in CC but shows a larger increase of ∼1.55 in BP from low to high disorder. Precision is largely maintained across bins, decreasing by at most 0.044 in BP and 0.043 in CC, and remaining within 0.045 of the low-disorder baseline in MF while recall decreases slightly with disorder by 0.036 in MF, 0.063 in BP, and 0.038 in CC indicating only mild sensitivity to highly disordered proteins. Together, these results suggest that PlasmoFP models learn relevant structure–function features from proteins with regions of intrinsic disorder, enabling function prediction even in scenarios where traditional sequence-based methods often fail.

**Table 4.** PlasmoFP performance (10% eFDR) with predicted intrinsically disordered proteins. Metrics are averaged per bin; Low, medium, and high disorder are (0.0-0.3), (0.3-0.6), (≥0.6) respectively across the entire protein sequence.

| | Disorder Content | # Proteins | $F_{\text{macro}}$ | $S_{\text{min}}$ | $\text{Precision}_{\text{macro}}$ | $\text{Recall}_{\text{macro}}$ | $ru$ | $mi$ |
| --- | --- | --- | --- | --- | --- | --- | --- | --- |
| MF | Low | 8911 | 0.901 | 2.087 | 0.910 | 0.892 | 1.892 | 0.880 |
|  | Medium | 1752 | 0.897 | 2.090 | 0.907 | 0.886 | 1.925 | 0.812 |
|  | High | 368 | 0.861 | 2.457 | 0.866 | 0.856 | 2.226 | 1.035 |
| BP | Low | 6905 | 0.853 | 3.679 | 0.917 | 0.798 | 3.626 | 0.622 |
|  | Medium | 1512 | 0.832 | 3.829 | 0.898 | 0.775 | 3.755 | 0.744 |
|  | High | 372 | 0.803 | 5.230 | 0.887 | 0.735 | 5.159 | 0.834 |
| CC | Low | 7466 | 0.880 | 1.147 | 0.939 | 0.828 | 1.132 | 0.182 |
|  | Medium | 1869 | 0.863 | 1.384 | 0.922 | 0.810 | 1.363 | 0.237 |
|  | High | 443 | 0.850 | 1.733 | 0.922 | 0.789 | 1.715 | 0.234 |

### PlasmoFP accurately predicts terms for experimentally validated proteins

We evaluated PlasmoFP performance on a subset of *P. falciparum* proteins held out from training. Two experimental studies and a bioinformatics analysis (*39-41*) recently characterized 353 RNA-associated proteins with annotations such as: RNA binding, mRNA splicing via spliceosome, RNA processing, translation, and nucleus. We masked existing annotations for this subset across all subontologies and used PlasmoFP models to predict GO terms. PlasmoFP BP 20% eFDR recovered key RNA-related BP terms for proteins with predictions greater than 3 nodes away from the root, including RNA processing, RNA metabolic process, regulation of gene expression, and RNA splicing **(Fig. S9)**. Predictions from the MF subontology were dominated by nucleic acid binding and purine ribonucleoside triphosphate binding, while the CC subontology highlighted intracellular membrane/non-membrane bound organelles and nuclear protein-containing complex. Proteins lacking deeper predictions were consistent with the applied eFDR control. Together, these results demonstrate that PlasmoFP generalizes effectively to previously unseen proteins, producing predictions that are consistent with experimental evidence.

### PlasmoFP models predict GO terms for partially annotated proteins

Given that PlasmoFP models were trained independently on each ontology, we evaluated their ability to predict meaningful GO terms for partially annotated SAR proteins using a cross-ontology imputation analysis (see Methods). This analysis specifically tests the model’s ability to complete missing annotations in one ontology using existing annotations from other subontologies across all SAR test sets as the background. At 20% eFDR, PlasmoFP achieves a mean imputation efficiency of 82.3% across 10,731 proteins, with ontology-specific efficiencies of 85% (95% CI 84–86) for MF, 83% (81.7–84.14) for BP, and 80.4% (79–81.7) for CC. Corresponding pairwise quality scores are 82.3% (95% CI 81.5–83.2) for MF, 80.2% (79.2–81.2) for BP, and 79.8% (78.6–81.0) for CC **(Fig. S10)**. Relative to a cross-protein shuffle control, pairwise quality improves by +7.99 (MF, paired-permutation p ≤ 0.001), +6.97 (BP, p ≤ 0.001), and +1.3 (CC, p=0.1512) percentage points (pp); versus a vocabulary-matched random baseline, gains are +7.91, +9.07, and +9.21 pp (p ≤ 0.001).

Results at 5% and 10% eFDR closely mirrored the 20% findings for MF and BP, while CC remained statistically indistinguishable from shuffle at all thresholds (20%: Δ=1.3 pp, p=0.15; 10%: Δ=0.67 pp, p=0.57; 5%: Δ=−2.44 pp, p=0.08) **(Fig. S10)**. The lack of separation from the shuffle control across all eFDRs suggests that PlasmoFP CC’s gap to the control is minimal, likely because lowering the eFDR reduces the CC prediction vocabulary and per-protein outputs (e.g., 114 terms and 1.6 predictions/protein at 20%; 98 terms and 1.4 at 5%). Therefore, this concentrates predictions on high-frequency, generic localization terms that co-annotate broadly with many MF/BP contexts. Shuffling generic terms across proteins produces similar cross-ontology co-occurrence patterns, compressing the difference to the model. In addition, CC–BP/MF coupling is intrinsically weaker in this evaluation, hence protein-level improvements over shuffle are muted even though performance remains above random. Together, these results demonstrate that PlasmoFP models can effectively predict meaningful, contextual GO terms for partially annotated proteins, extending their benefits from *Plasmodium* PUFs (no GO annotation) to proteins lacking annotation in one or two GO subontologies.

### New functional annotations for partially annotated and proteins of unknown function in *Plasmodium*

We utilized PlasmoFP models to predict GO terms for PUFs and partially annotated proteins (excluding multigene families) across 19 *Plasmodium* species **(Table S1)** over a range of eFDR thresholds. By integrating existing PlasmoDB proteome annotations with PlasmoFP predictions at 30% eFDR, we increased the fraction of proteins annotated with all three GO subontologies by an average of 35.3% and reduced the unannotated fraction by 25.3% across all species **(Fig. 3)**. Relaxing the eFDR threshold drove a steady increase in full annotation. By 10% eFDR, the unannotated fraction had fallen by a 73.2% average relative to baseline. Beyond 20% eFDR, the unannotated fraction plateaued. Meanwhile, the fully annotated set climbed by an additional 11.1% between 20% and 30% eFDR. The partially annotated category spans ∼32.9–57.8% at the existing baseline, peaks at 67.8% around 10% eFDR, and then declines as those proteins transition into the fully annotated pool at higher eFDR thresholds. Notably, PlasmoFP predictions elevated particularly poorly annotated species such as *P. cynomolgi*, *P. yoelii*, and *P. ovale wallikeri* to levels approaching the more richly annotated *P. falciparum* proteome. Even at a stringent 5% eFDR, these species achieve fully annotated fractions of 16.2–19.2%, a 90.5% average relative increase over the existing baseline. Conversely, *P. falciparum*, already at 42.3% fully annotated coverage, showed the smallest absolute gain, reflecting the comprehensive annotation coverage already provided by community efforts(*42*).

**Fig. 3.**
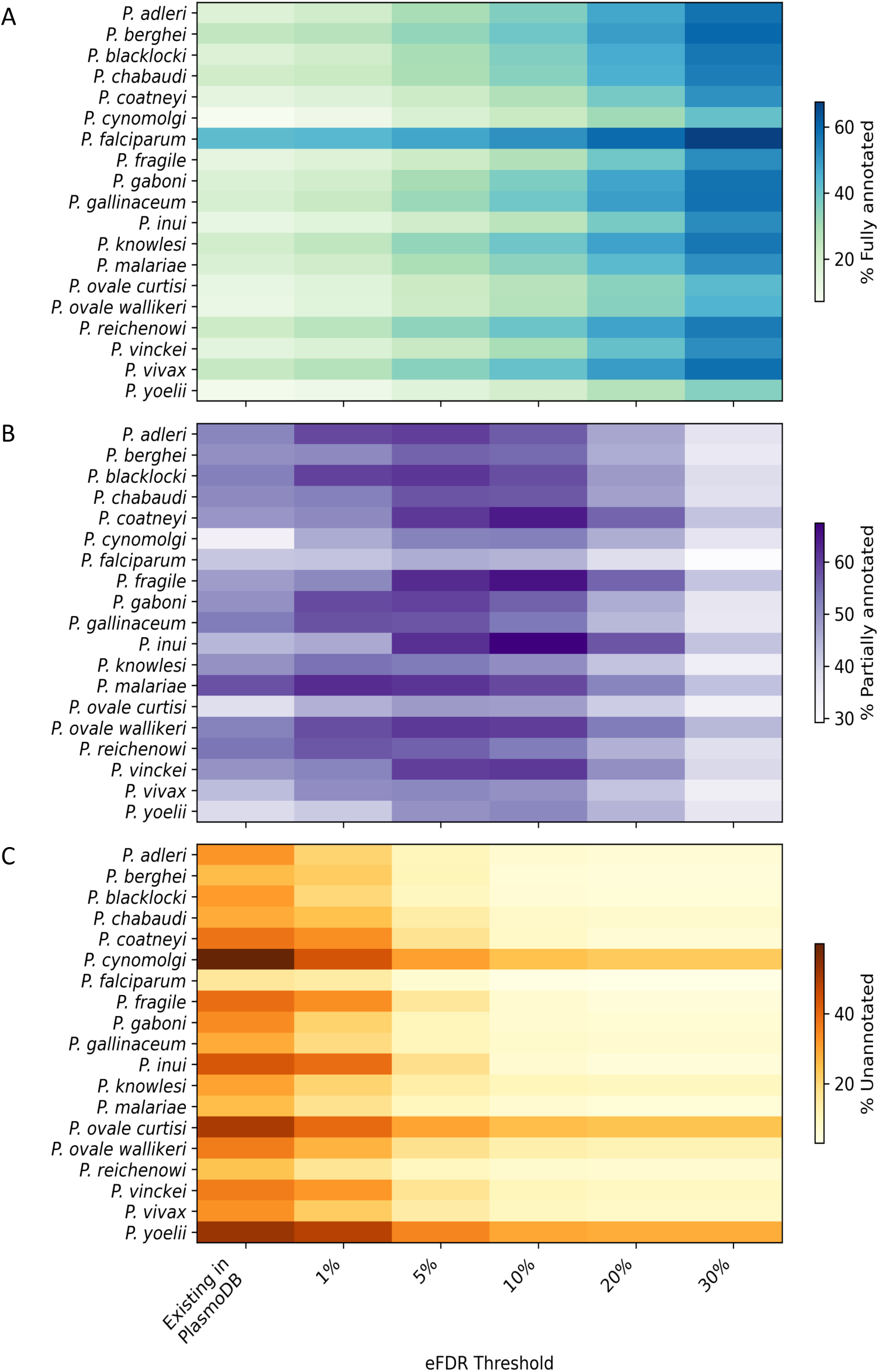
PlasmoFP models predict new functional annotations for *Plasmodium* proteins. (A) Fraction of proteins in each species which have annotations in all three GO subontologies after integrating PlasmoFP predictions with existing PlasmoDB annotations at specified eFDR thresholds. The “Existing in PlasmoDB” column refers to curated + IEA annotations found in the current GAF for the species from PlasmoDB. **(B)** Fraction of proteins in each species which have annotations in at least one GO subontology in the PlasmoFP + PlasmoDB dataset. **(C)** Fraction of proteins in each species which lack annotations in all three GO subontologies in the PlasmoFP + PlasmoDB dataset.

To test whether the PlasmoFP predictions are biologically plausible, we evaluated how novel GO terms relate to existing annotations and functional groupings. We first computed semantic-distance clusters using Resnik’s measure and then applied UMAP for dimensionality reduction(*43*). We found that newly predicted terms consistently clustered alongside existing annotations across all species and subontologies, indicating strong biological coherence when extant sequences are encoded into the model’s latent space, suggesting this featurization is useful and relevant in an AFP setting **(Fig. 4)**. Next, we grouped GO terms into functional clusters **(Fig. S11)**. In the BP subontology, the largest proportions of PlasmoFP BP predictions across partially annotated and PUF proteins fell into three clusters: (1) a “RNA Processing & Modification” cluster, (2) an “Amino Acid & Fatty Acid Metabolism” cluster, and (3) a “Regulation of Biological Processes” cluster **(Fig. S12)**. This suggests that PlasmoFP fills gaps in pathways that are potentially critical for parasite development. In the MF subontology, most predictions mapped to the “DNA/RNA Binding & Polymerase activity” cluster, followed by “Redox Activity” and “Hydrolases & Phosphatases” clusters **(Fig. S12)**. An observed enrichment of transporter-related predictions in both BP and MF subontologies is noteworthy, given recent reports on the challenges of characterizing membrane proteins with standard proteomic methods(*44*).

**Fig. 4.**
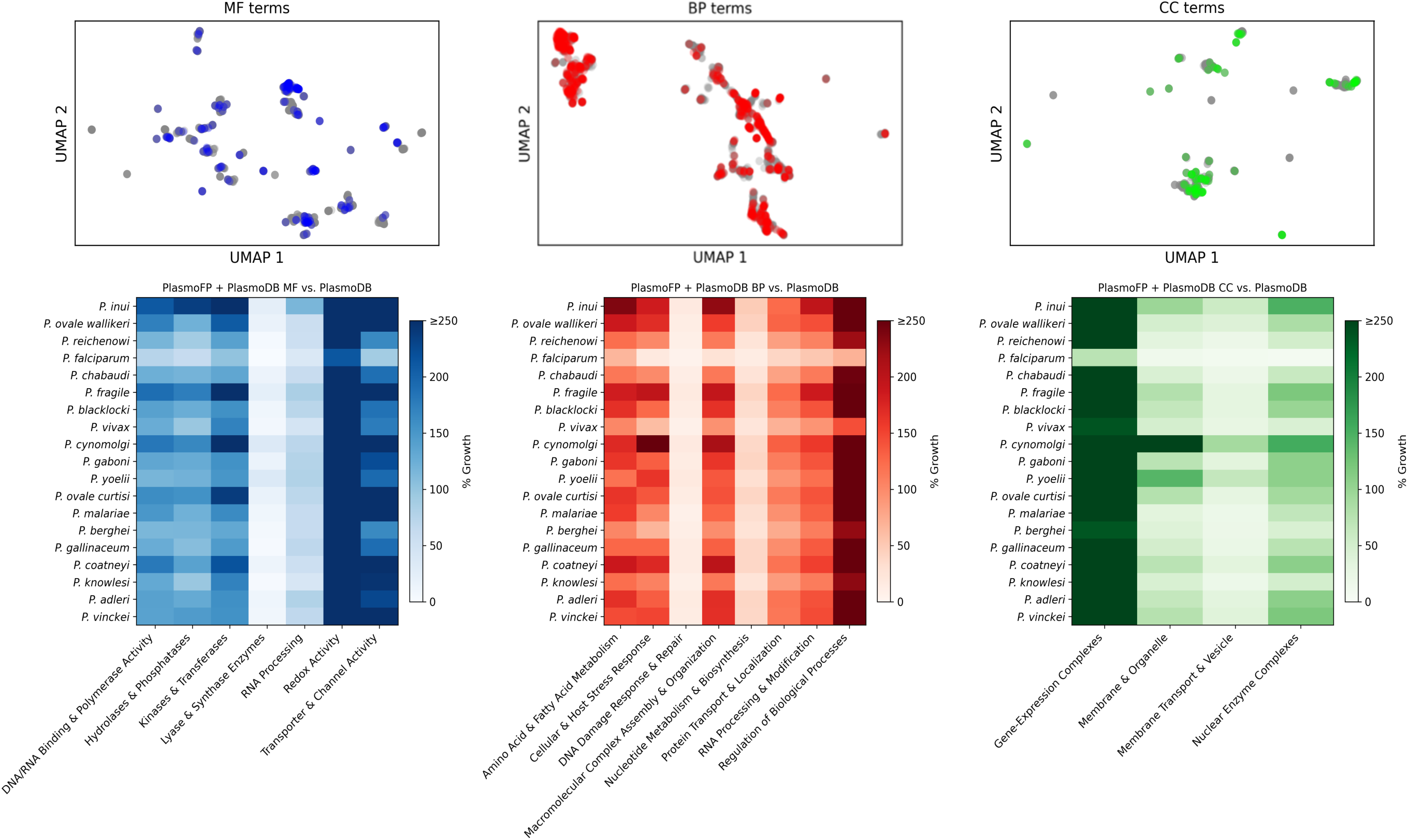
PlasmoFP predicted GO terms co-cluster with existing annotations and expand functional categories across subontologies. **(A)** UMAP projections of Resnik semantic-distance embeddings for GO terms in each subontology (MF: blue, BP: red, CC: green). Gray points are existing annotations; colored points terms predicted by PlasmoFP not found in the existing PlasmoDB annotation across all 19 *Plasmodium* species. **(B)** Heatmaps showing the increase in annotated proteins per functional cluster in the PlasmoFP + PlasmoDB set verses the PlasmoDB annotation.

When restricting our analysis to PUFs alone, the clustering distribution across subontologies closely mirrored that of the combined set of PUFs and partially annotated proteins, indicating that PUFs do not preferentially associate with any single functional cluster but rather distribute broadly across biological roles **(Fig. S12)**. This also implies that, since PUFs mirror the overall distribution, adding PlasmoFP predictions with existing annotations will not bias enrichment analyses or network reconstructions toward any particular functional category. Comparing the integrated dataset of PlasmoDB-existing BP annotations and PlasmoFP BP predictions against existing PlasmoDB BP annotations alone, we observed a 282.66% increase in proteins annotated within the “Regulation of Biological Processes” cluster **(Fig. 4)**. In the MF subontology, “Redox Activity” annotations grew by 400.6%, and in the CC subontology, the “Gene-Expression Complexes” cluster saw a 434.1% increase **(Fig. 4)**.

### PlasmoFP expands RNA-associated protein repertoire of *Plasmodium* species

*P. falciparum* has a limited transcription factor repertoire and shows a greater reliance on epigenetic and translation-level regulation of gene expression (*39, 45, 46*). However, predicting RNA-associated protein functions and RNA-binding remains a major challenge for AFP(*47*). To demonstrate the utility of PlasmoFP predictions, we explored whether they could recapitulate and extend defined RNA-associated protein repertoires in *P. falciparum* across the genus *Plasmodium*. Previous bioinformatic surveys catalogued 189 putative RNA-binding proteins spanning 13 families, including RRM (RNA recognition motif), KH (K homology), and zinc-finger domains(*40*); an HMM-based screen of 793 RNA-related domains annotated 18.1% of the proteome as RNA-associated(*39*); and proteomic profiling identified 898 RNA-dependent proteins, 545 of which had not been previously linked to RNA(*41*). In PlasmoDB, *P. falciparum* annotations catalog 920 RNA-associated proteins, comprising approximately 20.03% of the annotated proteome, while other species show substantially lower raw counts. PlasmoFP predictions in all functional categories added 121 new RNA-associated candidates in *P. falciparum*, refining the RNA-associated fraction of the annotated proteome to 19.90% from 20.03% **(Fig. 5A)**. This decrease indicates that, even as PlasmoFP predictions expand overall annotation coverage, the relative proportion of RNA-associated proteins to other functional groups of proteins remains unchanged. This stability also supports findings from previous experimental and computational surveys in *P. falciparum* (*39-41*).

**Fig. 5.**
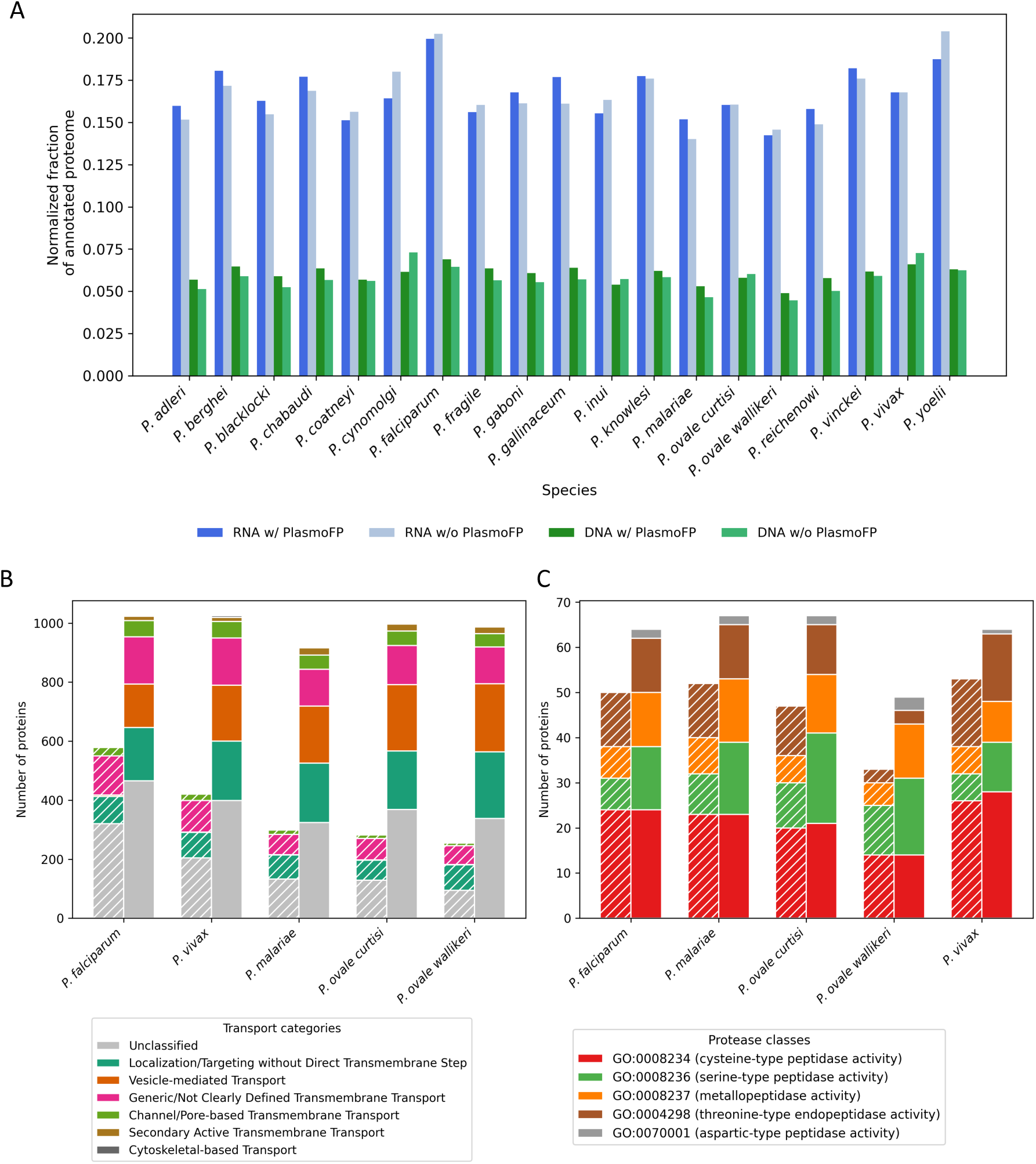
PlasmoFP predictions uncover new *Plasmodium* biology. **(A)** RNA- vs. DNA-associated proteome fractions in the *Plasmodium* genus. For each species, bars show the fraction of the annotated proteome labeled as RNA-associated (blue) or DNA-associated (green), before PlasmoFP integration (pale shading) and after adding PlasmoFP predictions to PlasmoDB annotations(solid shading). Normalized to each species’ total annotated proteome after integrating PlasmoFP 30% eFDR predictions. **(B)** Transport & localization counts across human-infecting *Plasmodium* species. Stacked bars for the five major human-infecting species show counts of proteins in Transport & Localization BP clusters mapped to TCDB-inspired categories. Proteins can belong to more than one cluster. Solid fill = PlasmoFP 30% eFDR predictions; hatched fill = existing PlasmoDB annotations. **(C)** Protease-like protein counts across human-infecting *Plasmodium* species. Stacked bars for the five major human-infecting species show numbers of proteins annotated as each protease class (cysteine, serine, metalloprotease, aspartic peptidase, threonine peptidase, and aspartic-type endopeptidase). Solid fill = PlasmoFP 20% eFDR predictions; hatched fill = existing PlasmoDB annotations.

Across the *Plasmodium* genus, PlasmoFP predictions uncovered considerable annotation gains, with the largest increases observed in the non-human primate species *P. cynomolgi* (+330 proteins), *P. inui* (+322 proteins), and *P. fragile* (+284 proteins) **(Fig. S13)**. Consistent with *P. falciparum*, despite these large additions, the normalized fraction of RNA-associated proteins in the annotated proteome remains stable at ∼15-20%, across all 19 species, changing at most by 1.65% in *P. yoelli* **(Fig. 5A)**. Additionally, we find the RNA-associated fraction of the PlasmoFP integrated proteome to be two-to three-fold higher than the DNA-associated fraction, demonstrating that PlasmoFP increases coverage without distorting functional balance across species. Collectively, these results suggest that a large RNA-associated proteome is a conserved, genus-wide feature, underscoring the importance of post-transcriptional control mechanisms across *Plasmodium*(*48*).

### PlasmoFP predictions expand existing functional classes in malaria parasites infecting humans

Proteases in *P. falciparum* are essential for parasite survival, mediating hemoglobin digestion (via aspartic, cysteine, and metalloproteases), and merozoite invasion/egress (via serine and metalloproteases)(*49*). However, ∼85% of *P. falciparum* proteases remain uncharacterized, and many protease-like proteins in other *Plasmodium* species lack any annotation(*49*). Protein transport and localization (protein sorting) constitutes another important aspect of malaria parasite biology, facilitating processes ranging from organelle targeting to nutrient uptake, but transport-related functions are under-annotated outside of *P. falciparum* and *P. vivax* (*44, 50, 51*). To assess how PlasmoFP fills annotation gaps in these two critical functional categories, protease-like enzymes and protein sorting, we investigated PlasmoFP 20% eFDR MF and 30% eFDR BP predictions across all *Plasmodium* species that primarily infect humans.

For proteases, we filtered PlasmoFP MF-predicted terms for aspartic, cysteine, serine and metallo-protease activities. PlasmoFP predictions added seven metalloproteases, five serine proteases, and two previously unannotated aspartic proteases in *P. falciparum*, boosting the total count of protease-like functions by 28% **(Fig. 5C)**. (Because our MF models could not be trained on threonine-type proteases due to insufficient training data for this term, that category showed no change.) When this approach was applied across all human-infecting *Plasmodium* species, the relative proportions of metalloproteases, serine, cysteine, and aspartic proteases remained similar, but species that are less well characterized realized larger absolute gains. For example, *P. vivax* and *P. ovale curtisi* gained 5 and 10 additional serine proteases, an 83% and 100% increase, respectively. Overall, predicted protease-like proteins increased by 34% on average across all human infecting *Plasmodium* species.

For transport/localization proteins, we took the subset of proteins found in the “Protein Transport & Localization” cluster, classified GO terms in this cluster into Transporter Classification Database(*52*)-inspired subclusters **(Table S2)** and counted the fraction of proteins in each subcluster **(Fig. 5B)**. The largest gain occurred in the “localization/targeting without direct transmembrane step” cluster, which rose from 1–5 proteins to 148–231 proteins across species. GO terms found in this cluster such as “establishment of protein localization to organelle”, “establishment of protein localization to membrane,” and “organelle localization by membrane tethering” that were previously under-annotated are now broadly represented in all five species. The “generic/not clearly defined transmembrane transport” cluster more than doubled in size on average while the “vesicle-mediated transport” cluster increased by up to 60 proteins in *P. ovale curtisi and P. ovale wallikeri*. This is noteworthy since vesicle-mediated transport is vital for protein export, nutrient uptake, and organelle maintenance(*53*). “Cytoskeletal-based transport” annotations, previously absent in the genus *Plasmodium*, appeared in all species with 2-6 proteins. While the “unclassified” cluster also increased with the PlasmoFP integrated dataset, the total number of unique proteins annotated with terms from the “Protein Transport & Localization” increased between 155-273 proteins across all species. Although *P. falciparum* and *P. vivax* remain highest in total transport annotations, *P. malariae*, *P. ovale curtisi*, and *P. ovale wallikeri* each gained ∼twofold more transport-related proteins.

## Discussion

Despite considerable progress in deep learning-based protein function prediction frameworks, no end-to-end methodologies exist to annotate uncharacterized proteins that have low sequence homology to well-characterized proteins within a phylogenetically defined subset of species. Existing models, such as DeepGOPlus and ProteInfer, are trained on large, phylogenetically agnostic datasets such as SwissProt, which consist primarily of well-characterized sequences from model organisms(*36, 37*). This approach often overlooks the unique characteristics of non-model organisms, where deviations from expected model performance are common due to remote homology instances. To address these challenges, we developed PlasmoFP, protein function prediction models tailored to a subset of non-model eukaryotic species found within the Stramenopiles–Alveolates-Rhizaria (SAR) eukaryotic supergroup that cause the global health disease malaria. We addressed challenges posed by both remote homology and non-model organism representation during training by leveraging protein structure representations using a fine-tuned structure-imbued protein language model, TM-Vec(*31*), and conditioning model learning on the phylogenetically relevant SAR structure-function relationships. By focusing on structure-function relationships, our approach to building a species-specific (*Plasmodium*) function prediction model can be generalized to non-model proteomes across the tree of life, provided their lineages have sufficient representation in annotation databases.

Although existing automated function prediction (AFP) techniques often provide confidence scores, few directly leverage uncertainty quantification to generate calibrated GO term prediction sets with explicit error-rate control. We found that by incorporating uncertainty using deep ensembles, we lowered the observed FDR in prediction sets across all subontologies in comparison to point models. Since PlasmoFP targets a subset of species rather than a broad spectrum, we also developed an FDR threshold-selection method that directly controls the false positive rate in prediction sets. Complementary structure-guided screens in *P. falciparum* using DALI on AlphaFold models have mapped folds to hundreds of PUFs(*35*); however, they do not produce calibrated GO term prediction sets or provide explicit error-rate control. PlasmoFP addresses this gap while retaining the benefits of structural signal. To evaluate PlasmoFP predictions against existing curated *Plasmodium* annotations, we developed a score to quantify biologically plausible GO term pairs across subontologies and found that PlasmoFP predictions generated annotation sets with co-occurrence patterns aligning closely with SwissProt proteins. PlasmoFP models were also able to maintain high performance on proteins with regions of intrinsic disorder, which are challenging to annotate using sequence-based deep learning techniques. This suggests that the intrinsic disorder content of a protein has little impact on learned structure-function relationships during model training. It also highlights the potential for developing a function prediction model for disordered proteins.

PlasmoFP performance was evaluated on a subset of experimentally validated *P. falciparum* proteins masked across all subontologies and produced predictions consistent with wet-lab experimental evidence. At the genus level, incorporating PlasmoFP predictions increased annotation completeness across all three GO subontologies and harmonized coverage in less-studied species. Semantic-distance clustering and functional cluster analyses indicate that the newly predicted terms are biologically plausible and fill gaps in protein repertories such as redox activity, transport, gene regulation. Together, these additions broaden and balance the functional landscape across the 19 *Plasmodium* species studied and provide a stronger and richer foundation for downstream multi-omics projects, molecular biology studies, and hypothesis generation and testing.

We also show that PlasmoFP predictions reveal expansions across functional classes which underpin *Plasmodium* biology. By expanding the RNA-associated proteome across the *Plasmodium* genus, PlasmoFP reinforces the importance of post-transcriptional control mechanisms across *Plasmodium* species. PlasmoFP predictions also expand protease and transport/localization annotations, two categories central to parasite invasion, egress, and nutrient acquisition. These additions close key annotation gaps in metabolic and trafficking pathways, offering new entry points for ongoing drug-target discovery efforts(*54*).

Collectively, these analyses show PlasmoFP captures both canonical and structurally conserved functional domains that traditional sequence-based annotation methods overlook. Importantly, PlasmoFP predictions for *Plasmodium* PUFs show no spurious enrichment in any single GO process, helping to prevent false-positive biases in downstream tasks such as GO term enrichment. The underlying framework used to train PlasmoFP is also compatible with additional information streams, including co-evolutionary features from MSAs(*55, 56*), per-residue sequence–structure embeddings, phylogenetic weighting, and calibrated uncertainty strategies. This positions PlasmoFP as a scalable, uncertainty-aware foundation for predicting annotation classes beyond GO across the genus *Plasmodium*.

In summary, approximately 20 years after the first malaria parasite species genomes were published(*4, 5, 57, 58*), we are now within sight of a comprehensive functional annotation. PlasmoFP advances this goal by closing critical annotation gaps and harmonizing annotation completeness between well-studied and lesser-studied malaria parasite species. Such protein function predictions help advance *Plasmodium* basic research and progress towards global malaria elimination.

## Materials and Methods

### PlasmoFP training and hyperparameter tuning

For each model in each ontology, we conducted a grid search over the following parameters: number of cascading hidden layers ([256], [256,128], [256,128,64], [256,128,64,32]), learning rate (0.01–0.00001), and number of epochs (10, 15, 20, 30, 40), using a standard binary cross-entropy loss function. During this grid search, the dropout parameter was fixed at 0.3 in the last layer before the output and a ReLU activation function was used for each layer. Each trained model was evaluated on the respective ontology validation set using the Fmax and Smin CAFA metrics, with the chosen model exhibiting the lowest scores for both (**Fig. S14**)(*20*).

To address potential class imbalance, we adjusted the binary cross-entropy loss function by weighting terms according to their depth and inverse frequency. Additionally, we explored alternative loss functions, including Focal, Jaccard, Hinge, and Asymmetric loss functions, but found that the unweighted binary cross-entropy consistently produced the lowest Fmax and Smin scores across all PlasmoFP models (**Fig. S14**) on the validation set.

### Model inference, evaluation, and output

For each GO subontology and protein–term pair, we infer a probability from our *k*-member deep ensemble by computing the median prediction minus its median absolute deviation (MAD). We then preform a parameter sweep over decision thresholds from 0.01 to 0.99, linearly spaced by 0.01, and for each GO term calculate the empirical FDR on the test set. If a term never appears in validation, we borrow the closest observed ancestor or descendant term (by the ontology graph) to estimate its FDR. This lets us map each binarization threshold to an observed FDR for every GO term. Finally, when we predict annotation sets for partially annotated or unknown proteins, we choose thresholds that guarantee a user-specified expected FDR. If a threshold is unavailable, we use the threshold corresponding to the maximum FDR ≤ target FDR.

We evaluated our models using standard CAFA metrics(*20*) reporting Fmax, Smin, remaining uncertainty (*ru*), and misinformation (*mi*) in “partial mode” analysis as outlined in(*20, 59*). For our intrinsic disorder evaluation, disorder was predicted using AIUPRED (*38*), with predicted disorder content for each protein calculated as the proportion of residues predicted to be disordered relative to the total protein length. We measured model performance using Fmax and Smin on bins of SAR test set sequences containing varying levels of predicted intrinsic disorder.

### Model uncertainty estimation

We approximate the posterior predictive with an ensemble

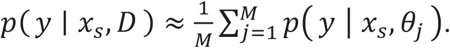

For each protein–GO pair we obtain *M* predictive outputs

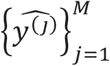

where *j* indexes the ensemble member produced by one of four approaches: (1) Deterministic baseline: *M*=1; (2) Monte Carlo Dropout (MCD): *M* stochastic forward passes with dropout enabled at inference; (3) Temporal ensemble: predictions from the last *k=5* checkpoints from model training (*M*=*k*); (4) Deep ensemble: *k* independently initialized models *(M=k)* on different folds of the training data. We compute an uncertainty-adjusted probability by taking the central estimate (ensemble mean or median of the softmax outputs) minus a dispersion measure (MAD or standard deviation). Models are trained on the full training set except for the deep ensemble method where each *k* member is trained independently with a distinct initialization.

### Training baseline models

*BLAST:* For each GO sub-ontology, protein sequence databases were built with makeblastdb (BLAST+ v2.15.0) from the SAR train set, SAR test set, and UniProtKB/SwissProt (accessed August 29, 2025). We performed BLASTp searches with an E-value threshold of 1×10^-5^ and returned up to 100 target sequences per query. Searches were run in two phases: first, testing sequences against the Swiss-Prot database, and second, testing sequences against the SAR training sequences. Baseline predictions were produced via annotation transfer from significant BLAST hits, using percent identity rescaled to a 0–1 confidence score. For each query, GO terms were gathered from all significant hits within the corresponding sub-ontology. When multiple hits supported the same GO term, we kept the maximum confidence score.

*FoldSeek:* Structural databases for the SAR train and test sets were created with FoldSeek’s createdb, using ProstT5 model weights to convert amino-acid sequences into 3Di structural alphabet representations. We then ran easy-search with an E-value filter of 1×10⁻⁵ and returned up to 100 sequences per query, searching both the Swiss-Prot structural database (AlphaFold DB representations) and the SAR training-set structural databases. Confidence scores for GO terms were derived by applying a sigmoid transformation to the bit scores of significant hits, scaled by the global standard deviation of all bit scores (with a minimum scale of 10.0 to avoid overly steep curves). As with BLAST, when multiple hits mapped to the same GO term, we retained the maximum confidence value.

*SAR CNN:* The SAR CNN models were trained and evaluated using the train and validation sets described earlier, with a one-dimensional convolutional neural network (CNN) architecture. The CNN architecture consists of two one-dimensional convolutional layers with kernel size 8 and padding of 4 to preserve sequence length. The first convolutional layer takes 21 input channels (corresponding to the one-hot encoded amino acids), while the number of output channels for both layers was optimized during hyperparameter tuning. Each convolutional layer is followed by ReLU activation functions and max-pooling operations with kernel size 2, resulting in a 4-fold reduction in sequence length. After the convolutional layers, the output is flattened and passed through two fully connected layers with dropout regularization (dropout rate optimized between 0.1-0.5). The architecture and training hyperparameters were optimized using Optuna with 20 trials per ontology. The optimization search space included: first convolutional layer output channels (32-512), second convolutional layer output channels (32-512), fully connected layer size (128-1,024), dropout rate (0.1-0.5), and learning rate (1×10^-5^ to 1×10^-2^ on log scale). Each trial was trained for 10 epochs, and the configuration maximizing Fmax on the validation set was selected for final model training. Final models were trained for 20 epochs using the Adam optimizer with binary cross-entropy. Training was performed with batch size 32, and sequences were processed in batches of 1,000.

*CAFA-CNN based on DeepGOPlus:* The CAFA-CNN models employed identical architecture and training procedures as the SAR CNN models but utilized the unified CAFA5 protein dataset from the CAFA5 competition [https://www.kaggle.com/competitions/cafa-5-protein-function-prediction]. The dataset was filtered in two stages: (1) retain only proteins with target ontology annotations (2) include GO terms present in the corresponding SAR training set vocabulary. Optuna hyperparameter optimization was performed with 20 trials on a 5% validation split randomly selected from the training data. The same optimization search space was used (convolutional channels 32-512, fully connected size 128-1,024, dropout 0.1-0.5, learning rate 1×10^-5^ to 1×10^-2^), with each trial trained for 10 epochs to maximize Fmax.

Final models were trained on the complete dataset using optimal hyperparameters for 20 epochs, maintaining the same technical specifications as SAR CNN (Adam optimizer, binary cross-entropy loss, batch size 32).

*CAFA-TM-Vec:* The CAFA-TM-Vec models utilized the same CAFA5 dataset as CAFA-CNN but employed the PlasmoFP neural network architecture instead of convolutional layers. Input consisted of 512-dimensional TM-Vec protein embeddings rather than one-hot encoded sequences. Hyperparameter optimization used Optuna with 50 trials on a 90/10 train-validation split, optimizing four predefined architectures ([256], [256,128], [256,128,64], [256,128,64,32]), five learning rates (0.01 to 0.00001), five epoch counts (10-40), and dropout rates (0.1-0.5). Models were trained with batch size 256 using Adam optimizer and binary cross-entropy loss, with final models trained on the complete dataset using optimal hyperparameters.

### Pairwise score

We assessed PlasmoFP’s ability to predict missing GO annotations in one ontology using existing annotations from the other two subontologies via a cross-ontology imputation analysis. For each protein annotated in ≥1 of the three subontologies, we masked its target-ontology annotations, generated PlasmoFP predictions for the target ontology from sequence, and then evaluated predicted versus real (masked) annotations against the protein’s existing cross-ontology annotations.

Term coherence was quantified with a pairwise quality score:

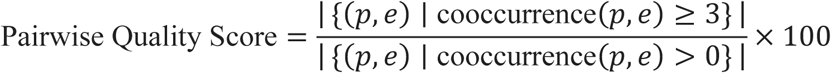

where *(p,e)* pairs each predicted target-ontology term *p* with each existing cross-ontology term *e* for the same protein. Co-occurrence counts were taken from the SwissProt unpropagated co-occurrence matrix (28,505 terms; \∼6.9M non-zero entries). Pairs with count ≥3 were counted as “good”; pairs with count >0 were “defined.” We applied deepest-term filtering to target-ontology predictions, and we applied the same filtering to ground truth and baselines to reduce parent–child artifacts before scoring.

Imputation efficiency was defined as:

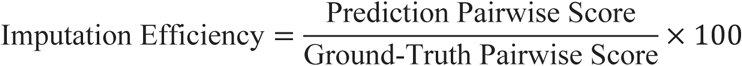

where the ground-truth score is computed identically after substituting the real (masked) target-ontology annotations for predictions. Pairwise scores (and the efficiencies derived from them) were computed as ratios of sums across proteins—i.e., we first sum “good” and “defined” pairs over proteins and then take their ratio. Analyses were run at eFDR thresholds of 20%, 10%, and 5%. To contextualize performance, we used two baselines evaluated with the same filtering pipeline: (1) a cross-protein shuffle, a uniform permutation (without replacement) that redistributes each model-predicted term set across proteins (testing protein-specific assignment while preserving term frequencies and set sizes); and (2) a pure random control that samples terms uniformly from the target-ontology prediction vocabulary, matching each protein’s predicted set size. Overall efficiency is reported as the unweighted mean of ontology-specific efficiencies.

For the four scores (prediction, ground-truth, shuffle, random), we computed 95% CIs via a nonparametric bootstrap over proteins, resampling proteins with replacement and recomputing the ratio-of-sums score within each replicate; intervals are percentile-based. For efficiencies (ratios of scores), we used a paired bootstrap over proteins (B=5,000), resampling the same indices for numerator (prediction, shuffle, or random) and denominator (ground truth).

*Significance testing:* To assess model-versus-control differences in pairwise score, we performed a two-sided paired permutation test across proteins (N=10,000 permutations). In each permutation, we randomly swapped (per protein) the two conditions being compared (e.g., model vs shuffle), recomputed the aggregate difference in pairwise score (ratio-of-summed “good” to “defined” pairs), and formed a null distribution. P-values used add-one smoothing,

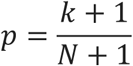

where *k* is the number of permutations whose absolute aggregate difference in pairwise score (computed as the ratio of summed “good” to summed “defined” pairs across proteins) was at least as large as the observed absolute difference, and *N* is the total number of permutations.

### Quality control

We performed quality control on the predicted annotations from all PlasmoFP models. Across all eFDR thresholds, we filtered out terms listed in the Gene Ontology Consortium’s *gocheck_do_not_annotate.obo* file. These are terms which are considered general and uninformative.

### GO term clustering by semantic distance

A GO term similarity matrix using Resnik for existing annotations combined with PlasmoFP predicted GO terms was calculated using biopython(*60*). An elbow plot was used to determine the optimal number of clusters for k-means clusters and generate word clouds using term names and manually labeled.

### Comparison with other models

We benchmarked PlasmoFP models against two state-of-the-art models: ProteInfer and DeepGOPlus(*36, 37*), selected for their demonstrated effectiveness in GO term prediction tasks. Both models were installed and configured locally in accordance with the instructions provided in respective repositories. Predictions were generated for the *Plasmodium* holdout set. For both ProteInfer and DeepGOPlus, we set the prediction threshold to zero, allowing us to retrieve all predicted probabilities associated with each GO term. This ensured that no potential predictions were excluded prior to post-processing. To enable a fair comparison, we applied the same evaluation criteria across all models. Using the *Plasmodium* holdout set predictions, we determined the threshold that optimized the Fmax metric for each model individually.

### Datasets

*Train, validation, and test set construction:* We assembled training, validation, test sets by collecting 907,145 proteins from the Uniprot release (2024_04): ‘(taxonomy_id:2698737) AND ((existence:3) OR (existence:2) OR (existence:1)) AND (length:[0 TO 1200]) NOT (taxonomy_id:5820) AND (active:true) AND (go:*) + ‘(taxonomy_id:5820) AND ((existence:3) OR (existence:2) OR (existence:1)) AND (length:[0 TO 1200]) AND (go_automatic:*) AND (active:true)’. Our criteria included proteins within the SAR supergroup, a protein existence score between 1-3, a total protein length ≤ 1200 amino acids, and exclusion of sequences with manually asserted GO terms in the genus *Plasmodium*. The dataset was partitioned by GO subontology into 716,735 MF, 573,293 BP, and 630,944 CC proteins.

Redundant sequences were eliminated using the MMseqs2 clustering function with parameters ‘–seq-id=0.9’ and ‘–cov=1’(*61*). Cluster representatives with the greatest number of GO terms were retained for maximum informational value. GO terms associated with each ontology were propagated to the respective root node using the ‘ancestors’ and ‘is_a’ queries from the QuickGO REST API for each GO term(*62*). Information Accretion for was calculated using the method described in and implemented via [https://github.com/claradepaolis/InformationAccretion], with release 2024-09-08 of the go-basic.obo(*59*).

Each ontology dataset was processed with MMseqs2 using ‘–seq-id=0.3’ and ‘–cov=1’ to form clusters of sequences sharing 30% similarity. We allocated whole clusters to the train, validation, or test sets using 80/10/10 partitions to prevent data leakage and removed terms with a frequency ≤ 50 across all sets to avoid training on low-frequency terms. This resulted in 289,930, 247,286, and 227,756 training sequences for the MF, BP, and CC subontologies, respectively; 29,188, 27,602, and 24,531 sequences for validation; and 33,343, 29,679, and 26,746 sequences for testing. The training label spaces comprised 744 MF terms, 1,548 BP terms, and 255 CC terms. For evaluation, GO terms in the validation and test sets were restricted to those present in the corresponding training label spaces.

*Plasmodium holdout set composition:* We collected 1,527 sequences from Uniprot (2024_04): ‘(taxonomy_id:5820) AND ((existence:3) OR (existence:2) OR (existence:1)) AND (length:[0 TO 1200]) AND (go_manual:*) AND (active:true)’. Our criteria included *Plasmodium* proteins possessing manually asserted terms and a total protein length ≤ 1200 amino acids. This dataset was also divided into three subsets, one for each GO ontology. GO terms for sequences in this set were not filtered to create the most representative version of the current annotation in the genus.

*Plasmodium PUFs and partially annotated set:* We processed *Plasmodium* genomes from Release 68 of PlasmoDB by parsing Annotated_Proteins.fasta files for each species and applying five sequential filters: (1) gene products longer than 1,200 amino acids were removed due to TM-Vec embedding limitations; (2) entries annotated as pseudogenes or partial genes were excluded based on description fields; (3) only primary-isoform transcripts ending in “.1”, “_1” or “_t1” were retained; (4) sequences belonging to large, variable multigene families were excluded by screening for the keywords “emp”, “msp”, “rifin”, “stevor”, “vir”, “fam-”, “pir”, “cir”, “yir” and “surf”; and (5) any gene annotated as part of the apicoplast genome was removed.

*P. falciparum experimentally validated RNA-associated proteins:* We excluded 353 proteins identified in (*41*) from all training datasets.

*TM-Vec embeddings*: We used the TM-Vec model to compute embeddings for sequences in all datasets as input for PlasmoFP and ALL-TM-Vec models(*31*). Each sequence was encoded using the ‘CATH’ model, resulting in a flattened embedding with a dimensionality of (1,512) per protein sequence.

## Acknowledgments

We thank Nathan Frey and Francisco Callejas-Hernandez for helpful discussions around project formulation, and Steven Sullivan, Mari Shiratori, and Kavya Phadke for editorial assistance. The NYU IT High Performance Computing resources, services, and staff expertise are also appreciated.

## Funding

This work was supported by the following research funds:

National Institute of General Medical Sciences, National Institutes of Health T32GM132037 (supported HS).

National Institute of Allergy and Infectious Diseases, National Institutes of Health U19AI089676 (JMC). The content is solely the responsibility of the authors and does not necessarily represent the official views of the National Institutes of Health.

Bloomberg Philanthropies as part of the support of Johns Hopkins Malaria Research Institute (JMC).

## Author contributions

Conceptualization: HRS, JMC, RB

Methodology: HRS, DB

Investigation: HRS, DB, OQ, ZW

Visualization: HRS

Supervision: JMC, RB

Writing - original draft: HRS, DB, OQ, ZW, RB, JMC

Writing - review & editing: HRS, DB, OQ, ZW, RB, JMC

Funding - JMC

## Data and materials availability

Training, validation and testing data splits for PlasmoFP can be found at https://doi.org/10.6084/m9.figshare.30100396.v1. PlasmoFP predictions for all 19 *Plasmodium* species can be explored at https://janecarltonlab.org/plasmofp. A Google Colab Jupyter notebook can be accessed at: http://tiny.cc/PlasmoFP. The source code for all key findings in this study are available on Github at https://github.com/harshstava/PlasmoFP_public. Raw dictionaries containing model predictions and the source code for the interactive Python webapp can be found at https://github.com/harshstava/PlasmoFP_Explorer.

