## Supplementary material for "PlasmoFP: leveraging deep learning to predict protein function of uncharacterized proteins across the malaria parasite genus": Fig. S

Harsh R. Srivastava *et al.*

**This PDF file includes:**

Figures S1 to S14

Tables S1 and Table S2

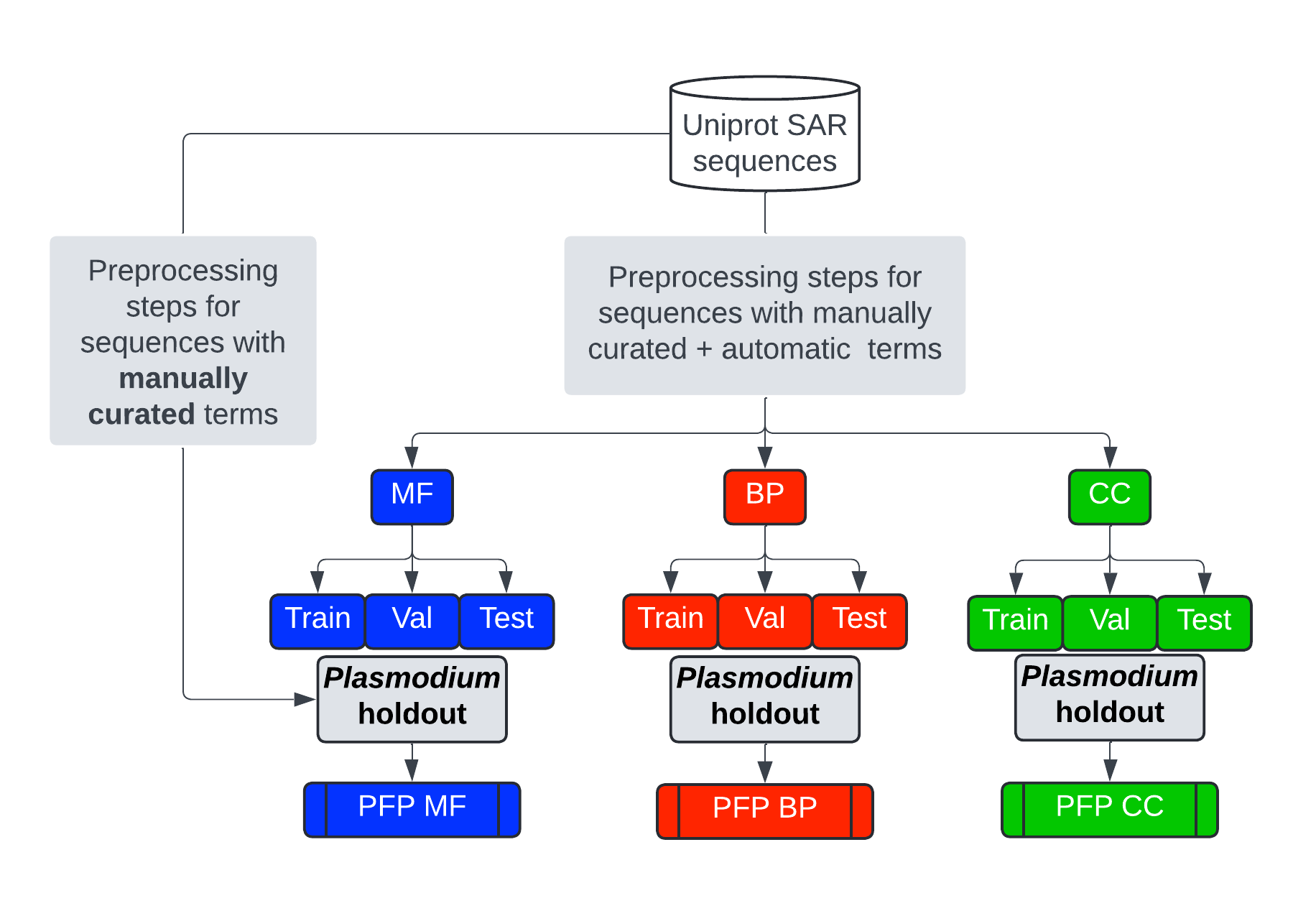

Figure S1. Data processing pipeline for PlasmoFP models. Processing pipeline regarding four datasets for each PlasmoFP model: train, val, test, and *Plasmodium* holdout.

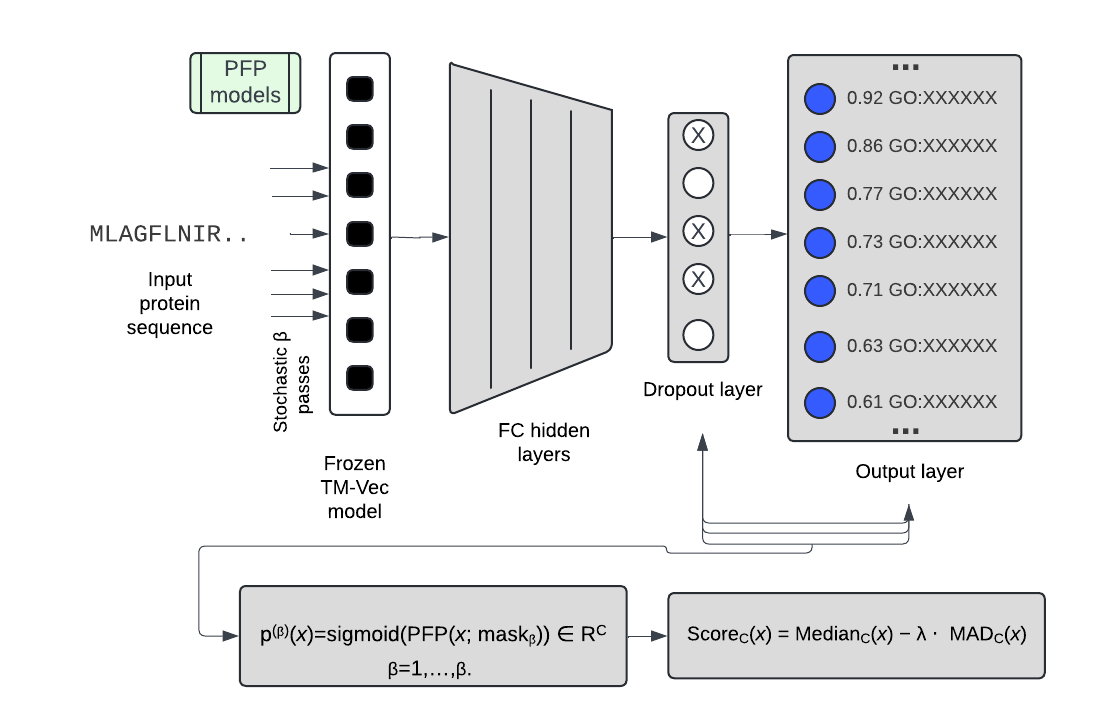

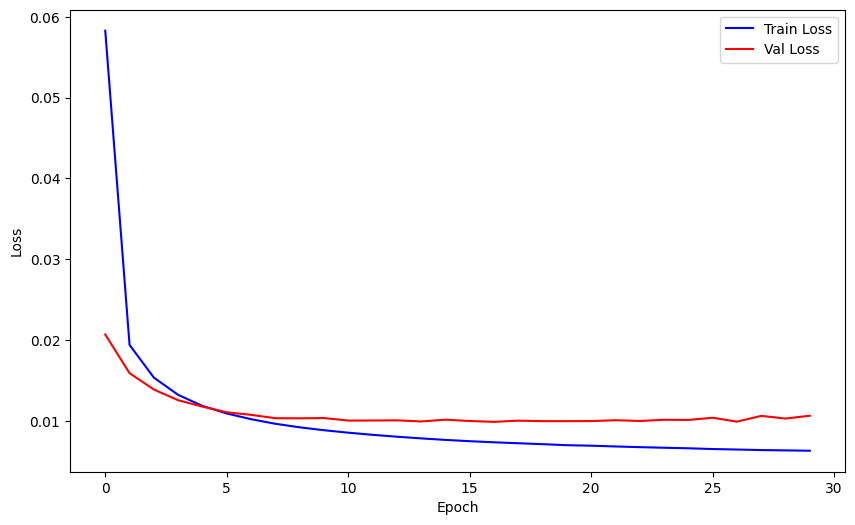

Final epoch

Final epoch - N

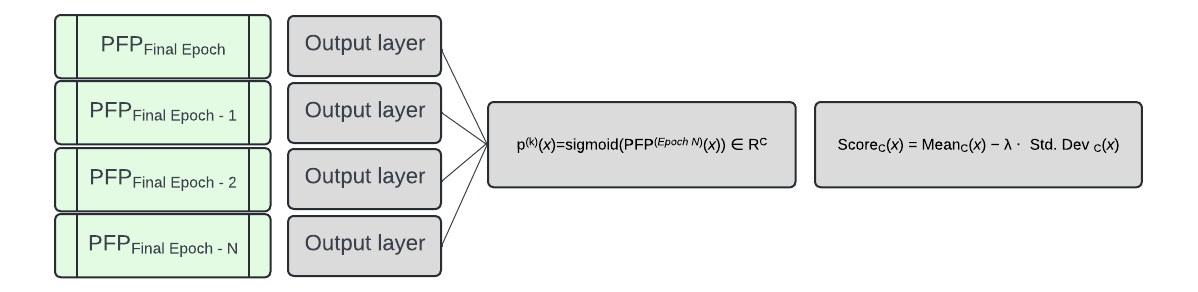

A

B

**Figure S2. Uncertainty-aware model schematics.** A) Schematic for Monte Carlo Dropout model trained using all available data. The dropout layer is turned on during model inference and multiple stochastic forward passes are made through the network. B) Schematic for a temporal ensemble which aggregates predictions from multiple checkpoints saved during model training on all available data.

A

B

C

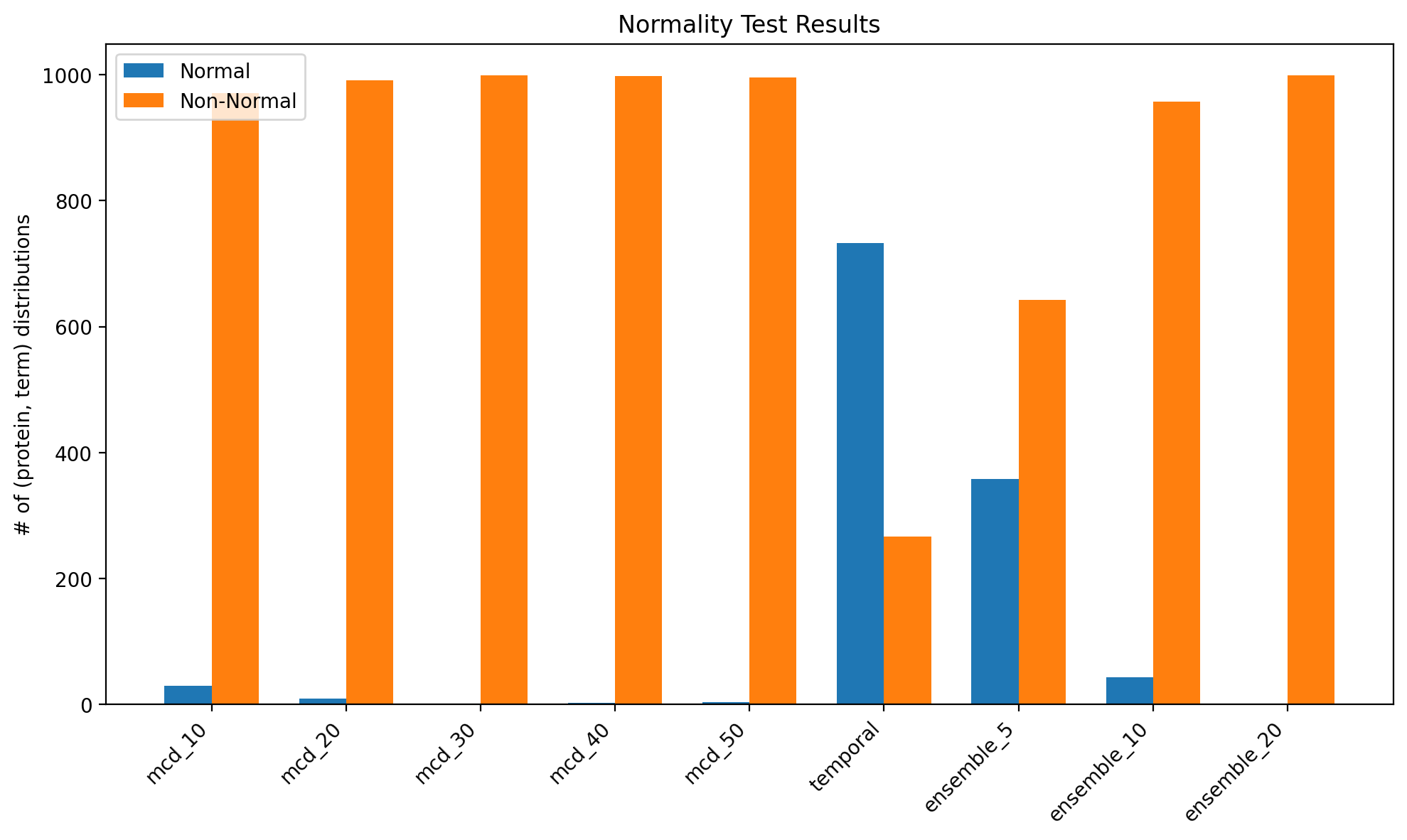

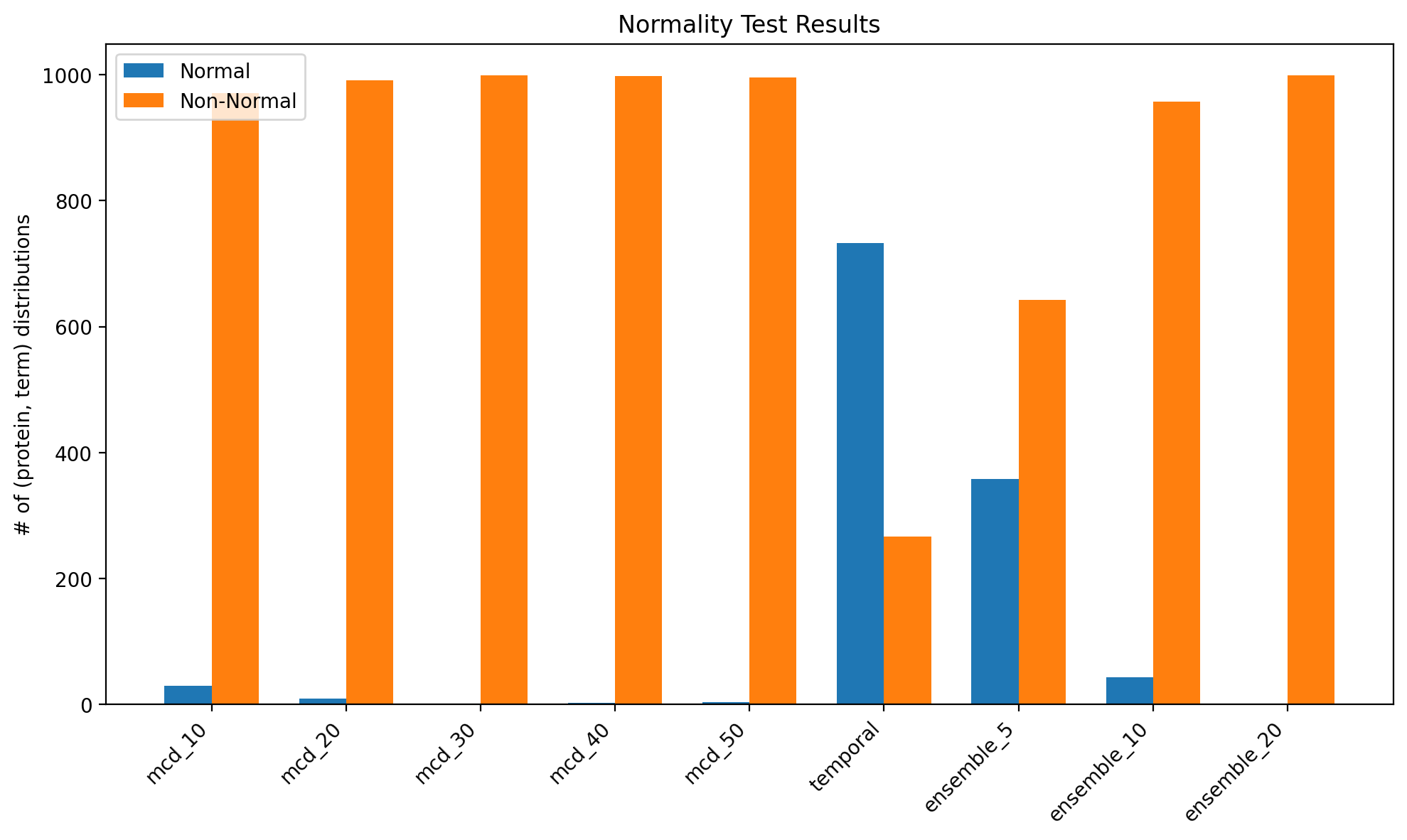

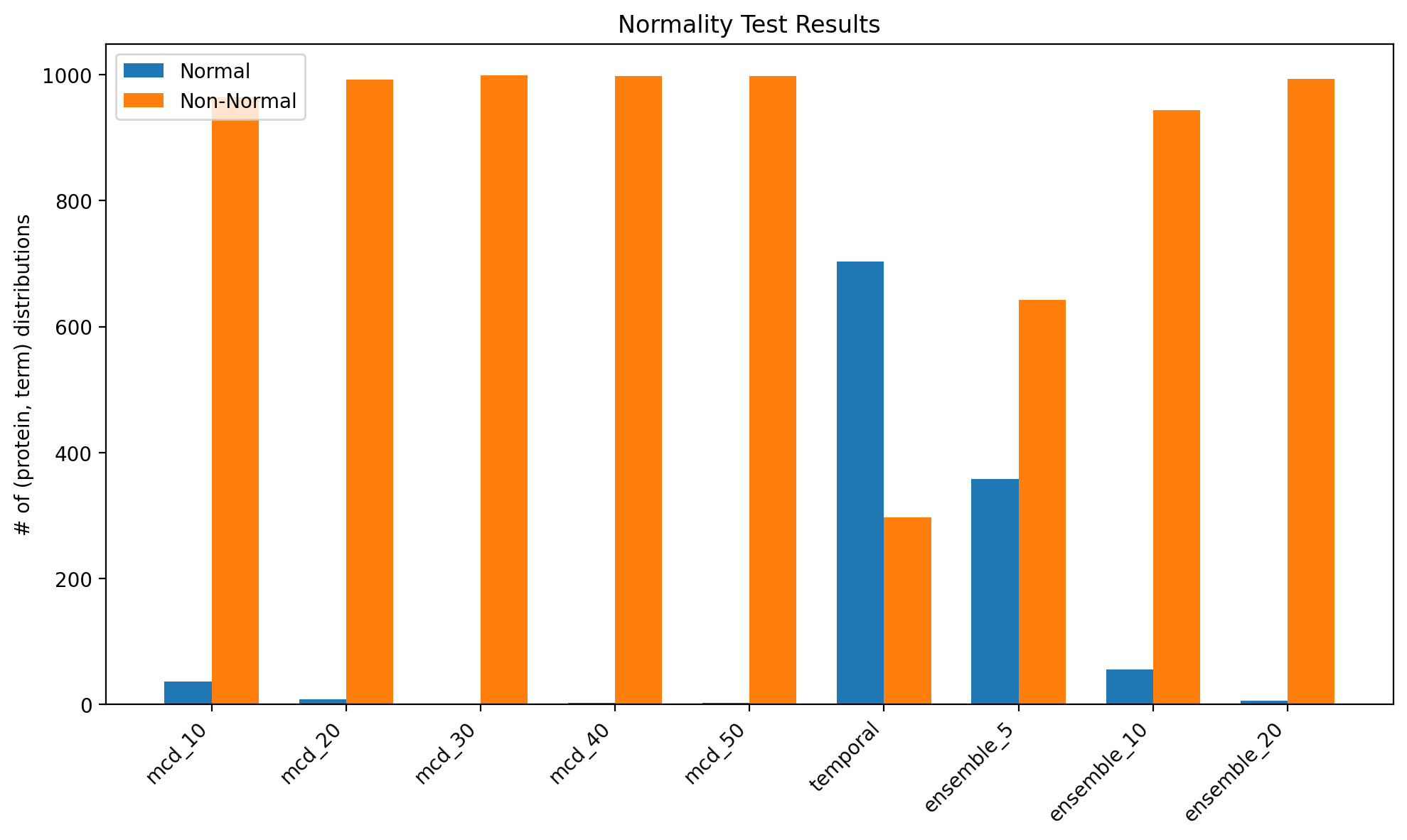

**Figure S3. Normality tests.** Shapiro–Wilk normality tests were performed on softmax-probability distributions for a random subset of protein–GO-term pairs, using each uncertainty estimation technique. Distributions with p > 0.05 are classified as ‘Normal.’ A) is the MF subontology, B) is the BP subontology, and C) is the CC subontology.

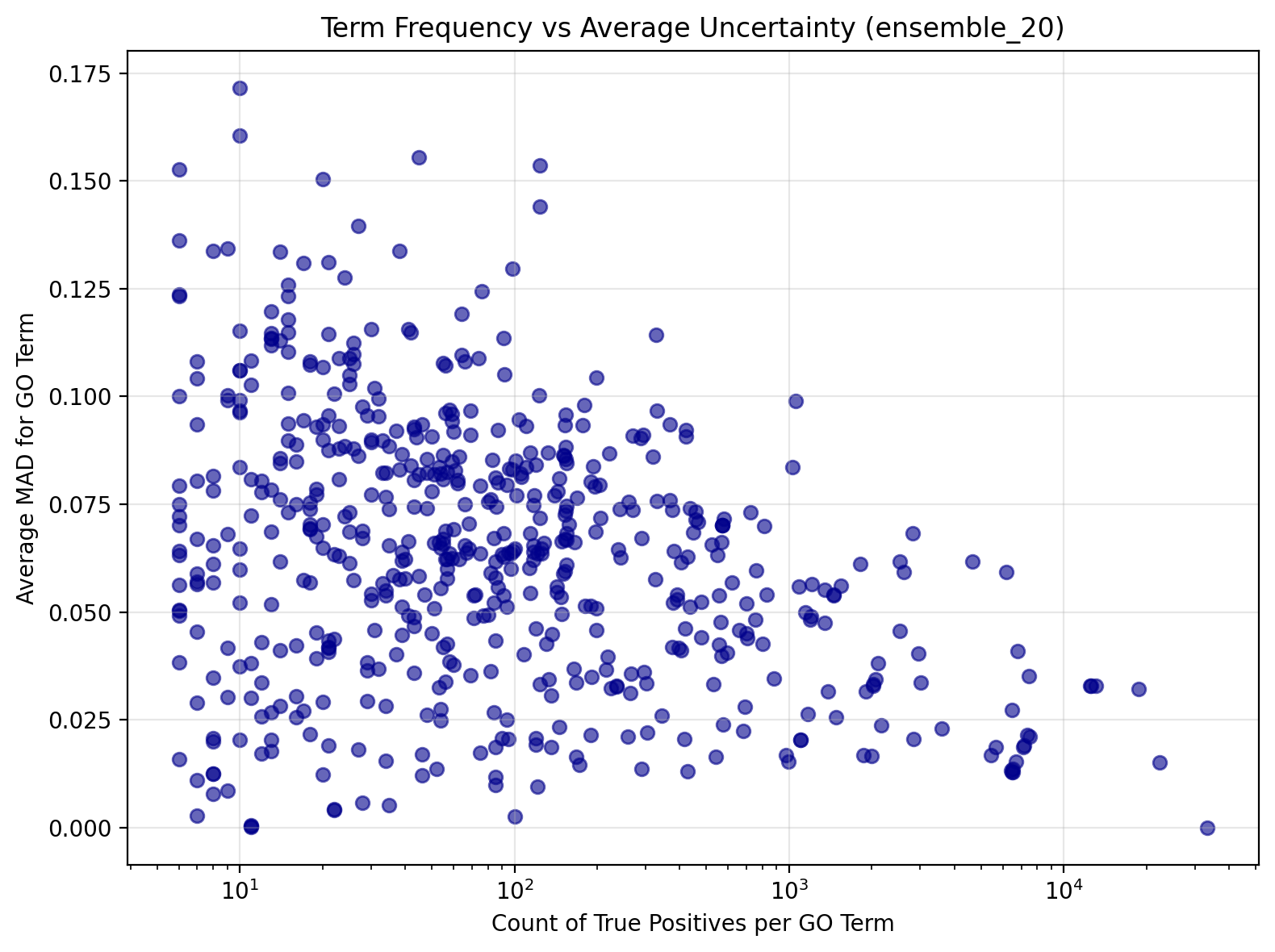

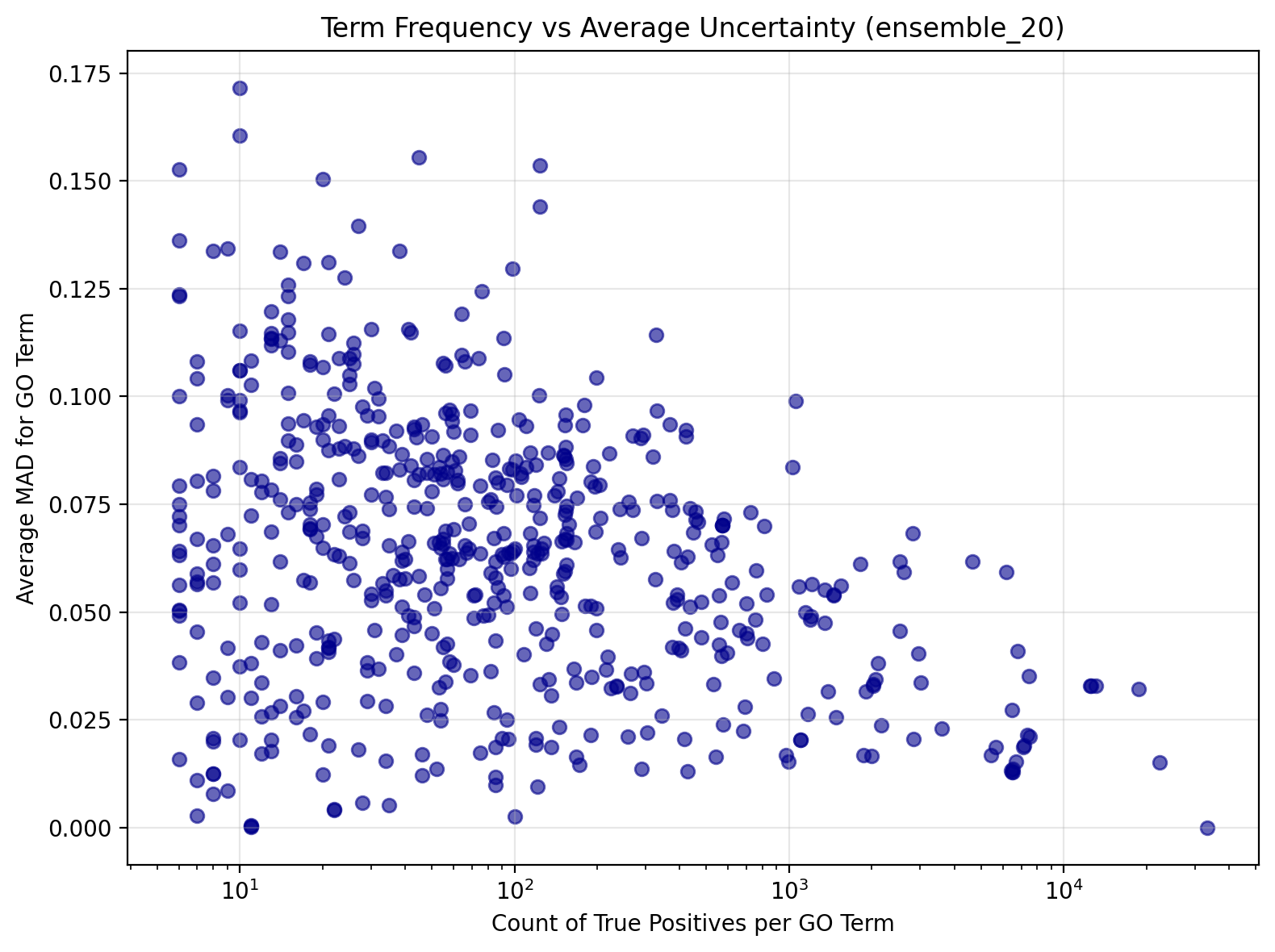

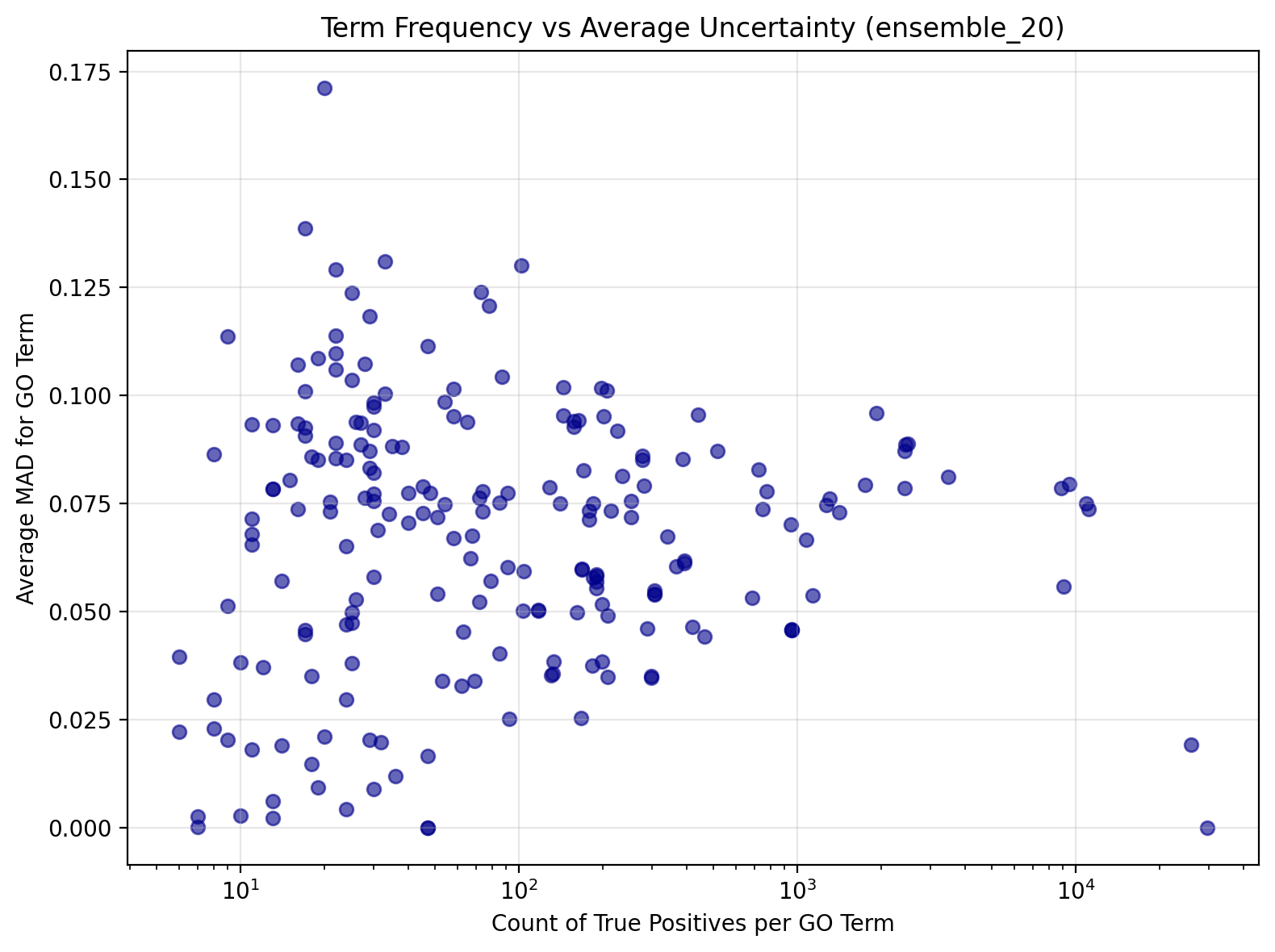

A

B

C

**Figure S4. Uncertainty as a function of true positives in the test set.** Scatterplots of the average epistemic uncertainty (MAD) for true‐positive predictions versus the count of true positives per GO term (x axes on a log₁₀ scale) in the test set, shown separately for: A) MF B) BP C) CC subontologies. Each point represents one GO term.

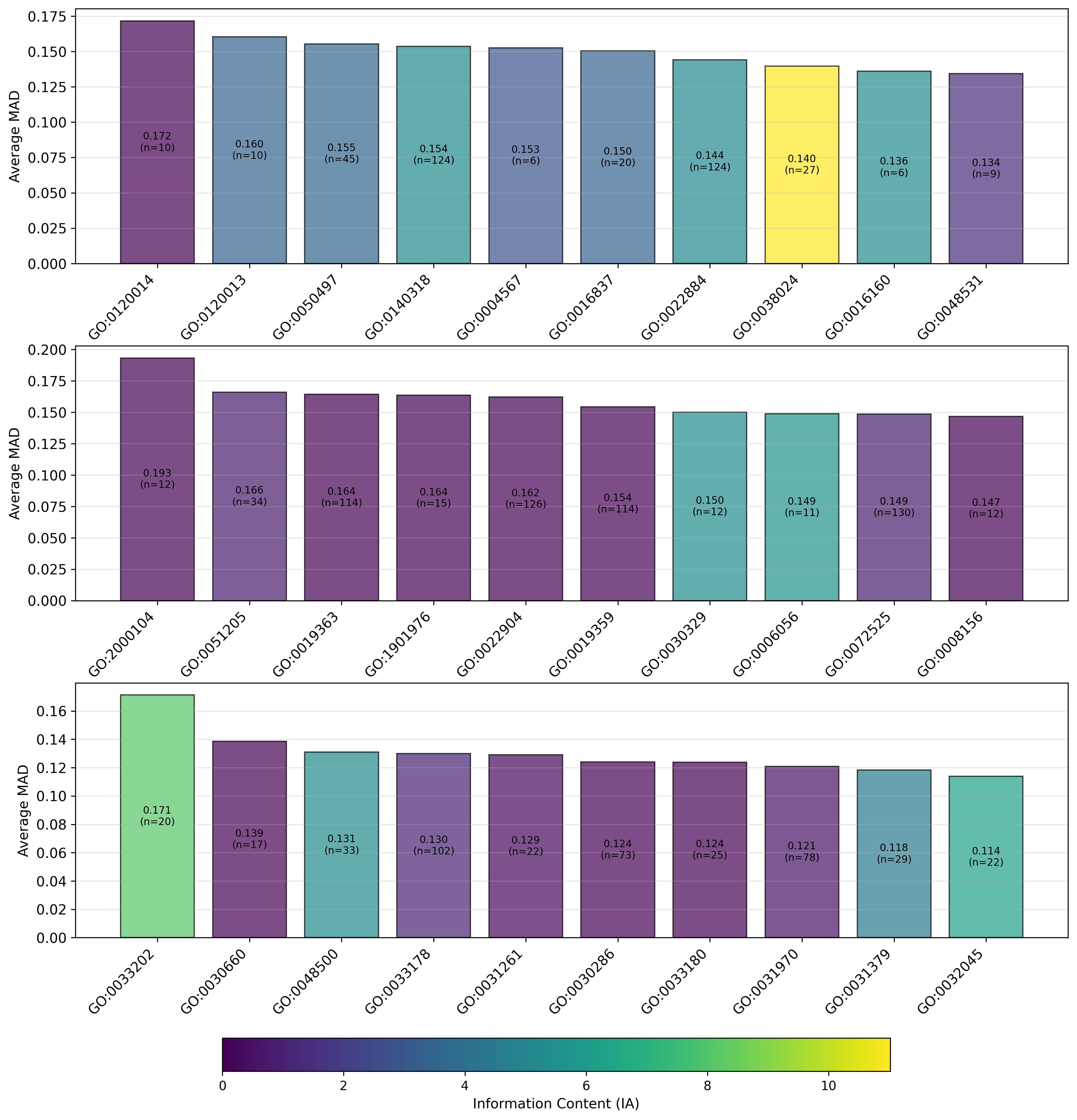

A

B

C

**Figure S5. Most uncertain terms for each PlasmoFP model.** A) MF, B) BP, and C) CC GO terms in the test set appearing ≥ 5 times, ranked by largest MAD.

A

B

C

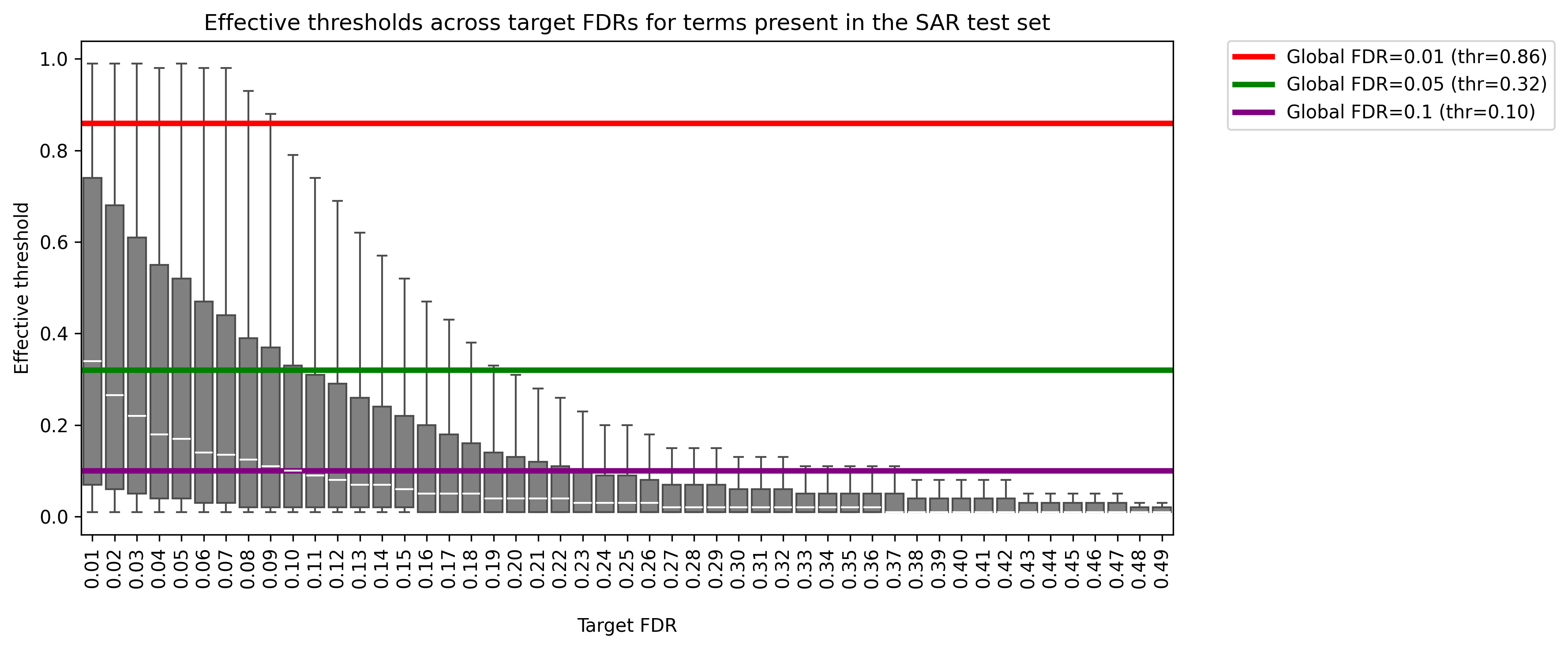

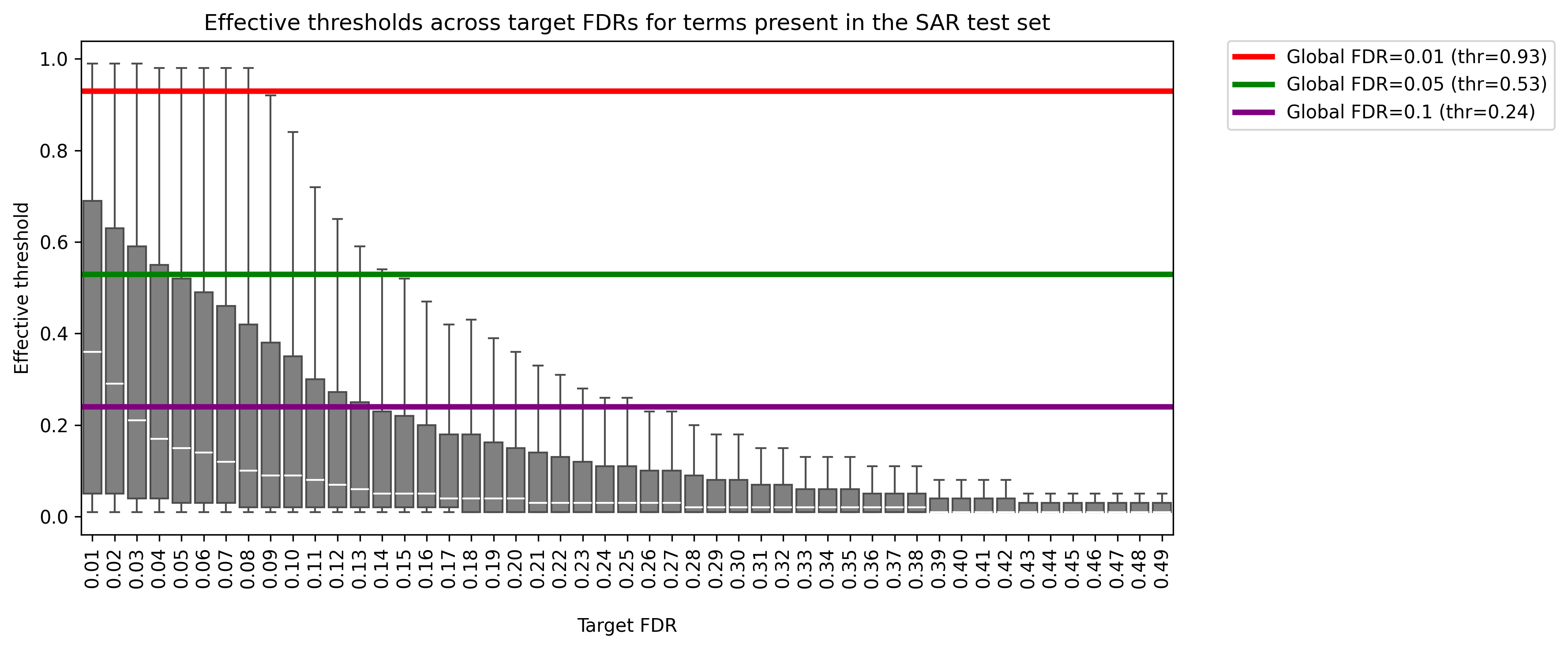

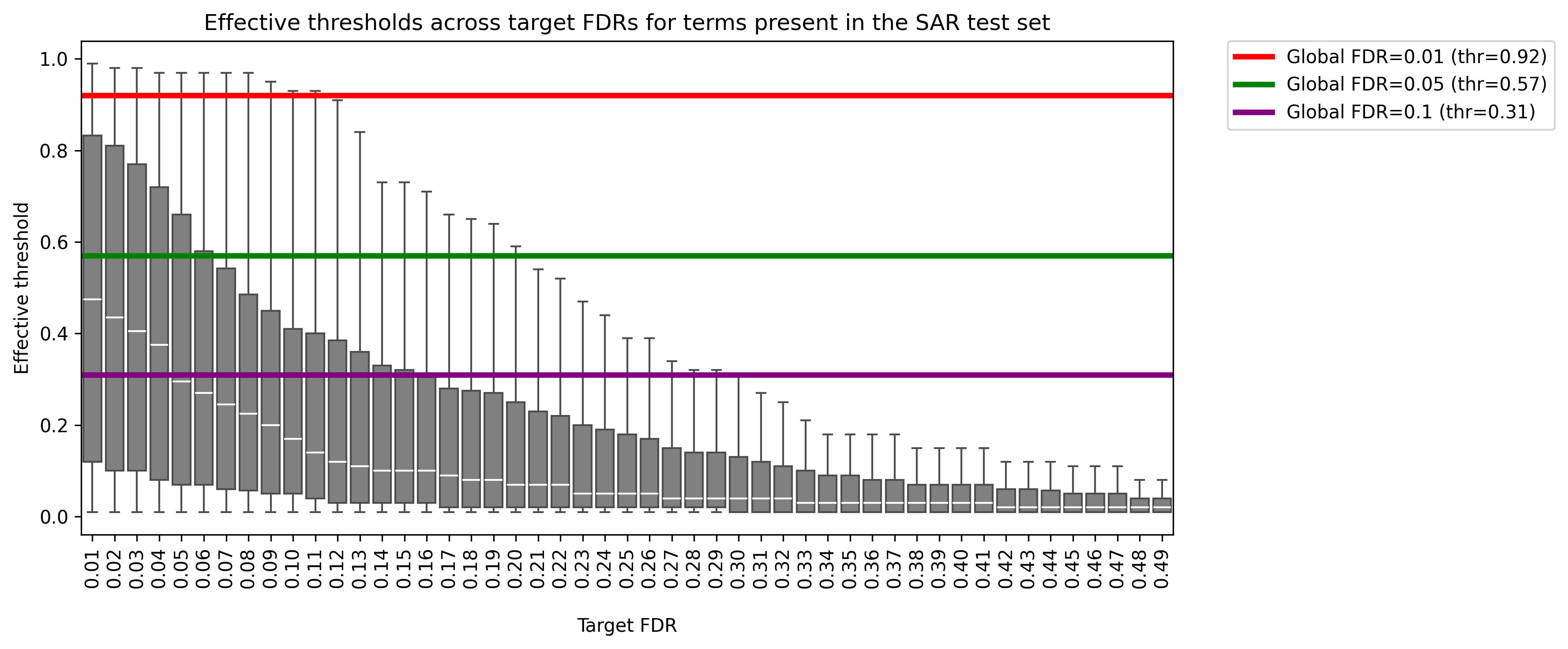

**Figure S6**. **Per-term thresholds for target FDR.** Boxplots show, for each target FDR on the x-axis, the distribution of “effective thresholds” (uncertainty adjusted softmax probabilities, median-MAD) required by individual GO terms to achieve that FDR, computed on the A) MF B) BP and C) CC SAR test sets.

A

B

C

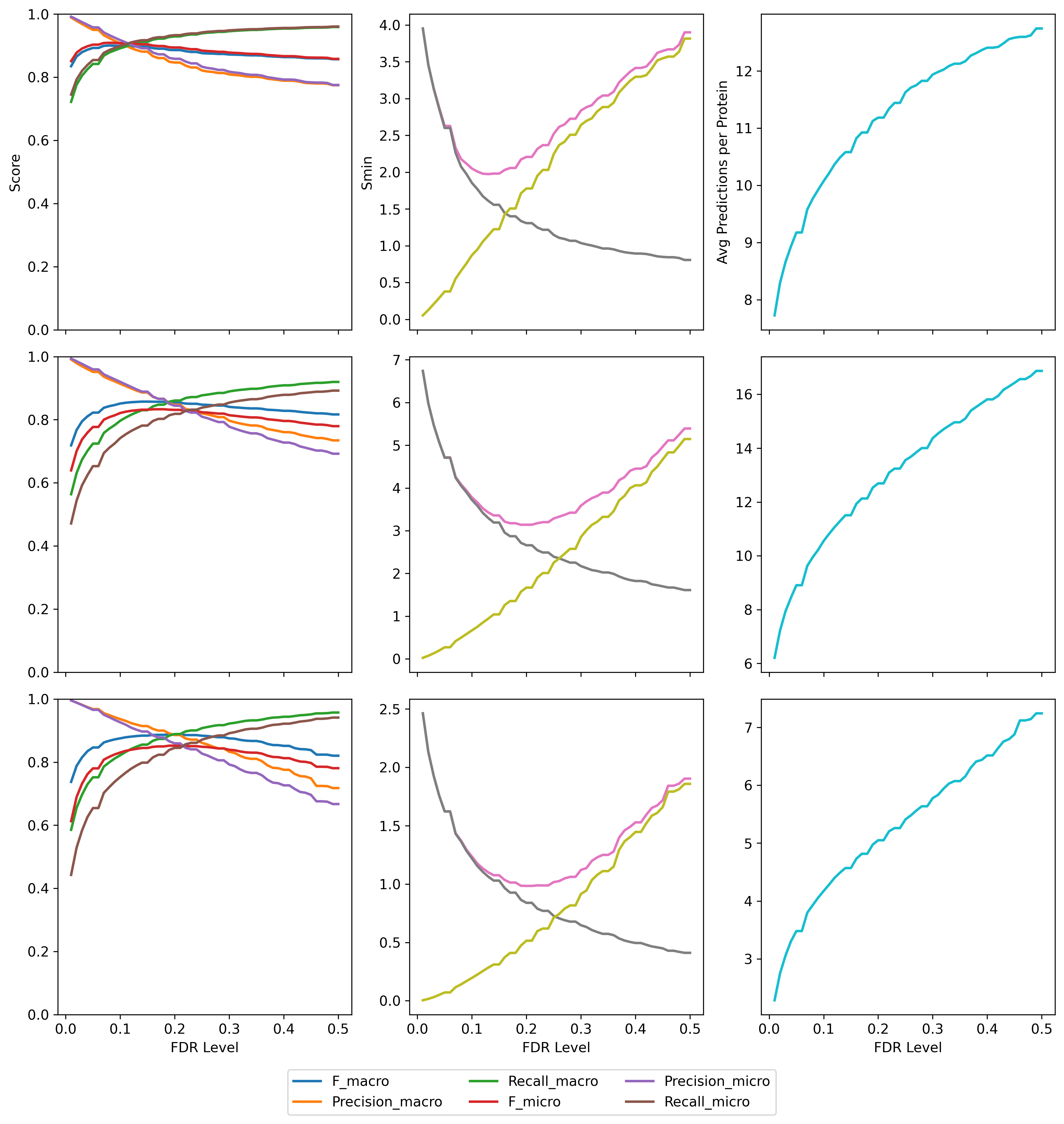

**Figure S7. Model performance using per-term FDR thresholds.** Performance curves shown using per-term FDR controlled thresholding on the A) MF B) BP and C) SAR test sets. Grey lines in S_min_ plots denote *remaining uncertainty* while yellow lines denote *misinformation* in the middle column.

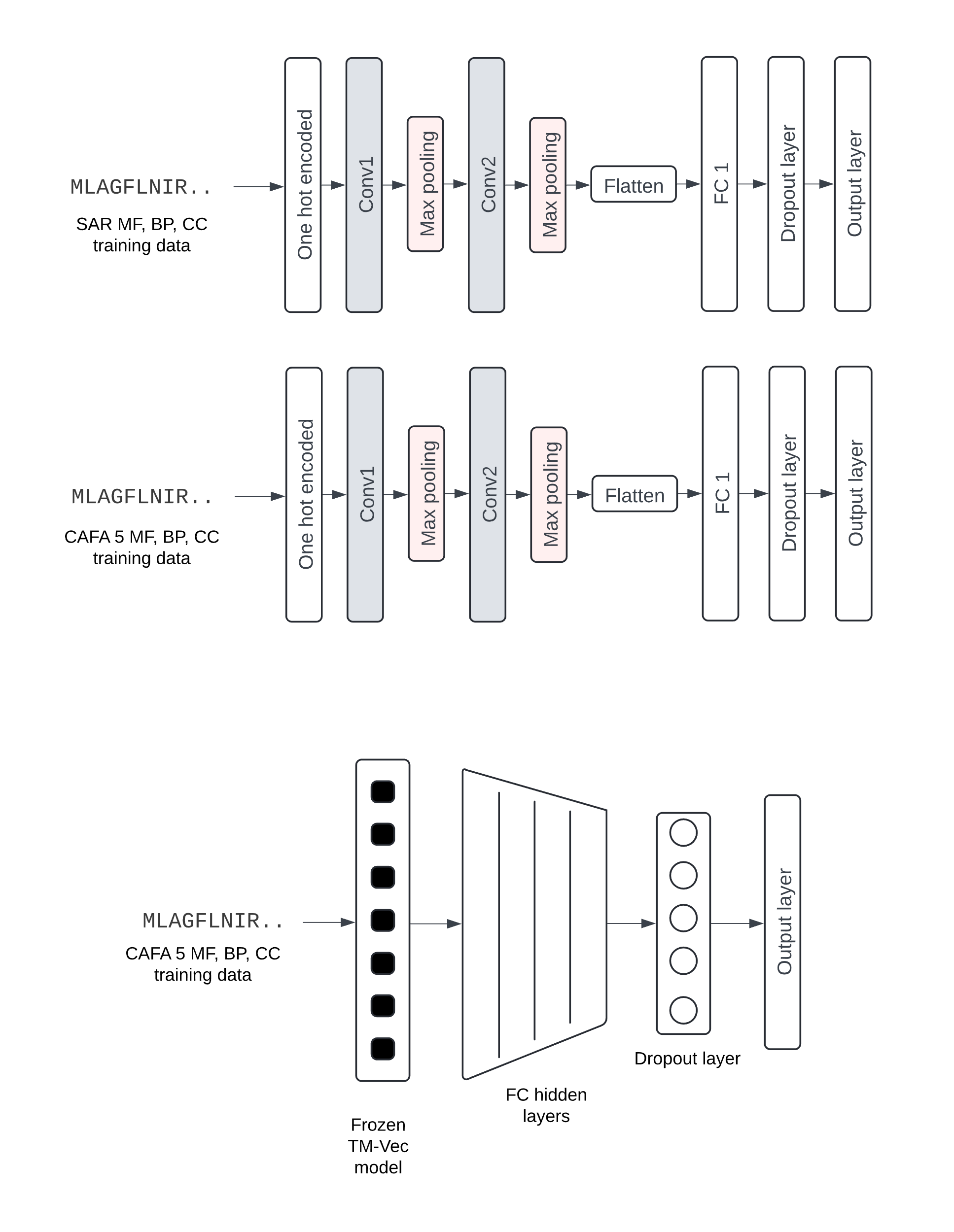

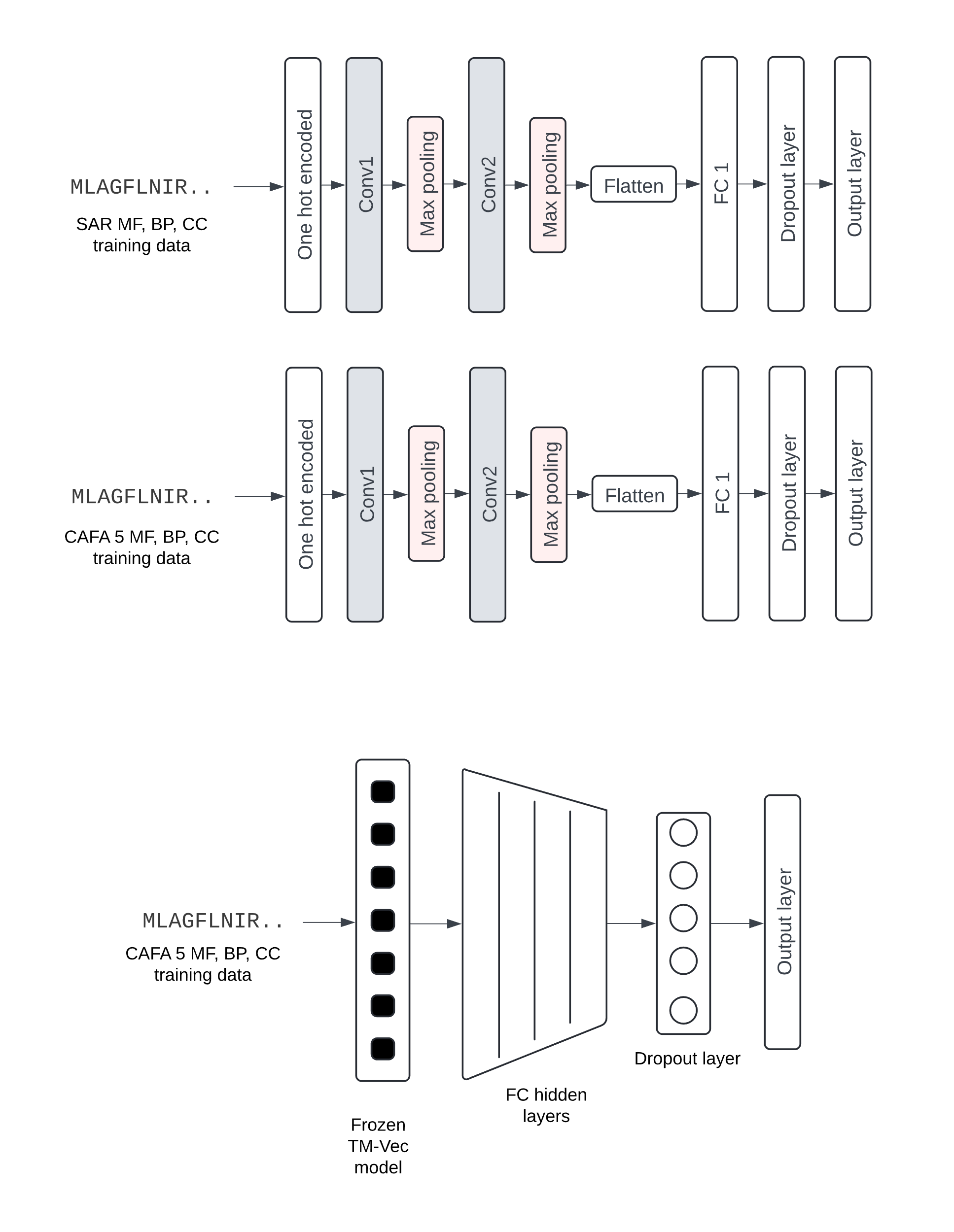

A

B

C

**Figure S8**. **Baseline-model architectures.** Model architectures for A) SAR-CNN MF, BP, CC, B) ALL_CNN MF, BP, CC, and C) ALL-TM-Vec MF, BP, and CC. Hyperparameters tuned for A and B include the number of filters in the first convolutional layer (conv1_out), the number of filters in the second convolutional layer (conv2_out), the size of the fully connected layer (fc_size), the learning rate (lr), and the number of training epochs (num_epochs) for each task (MF, BP, CC). Hyperparameters tuned for (C) include lr, # of layers before the dropout layer, the dropout rate (gamma) and num_epochs.

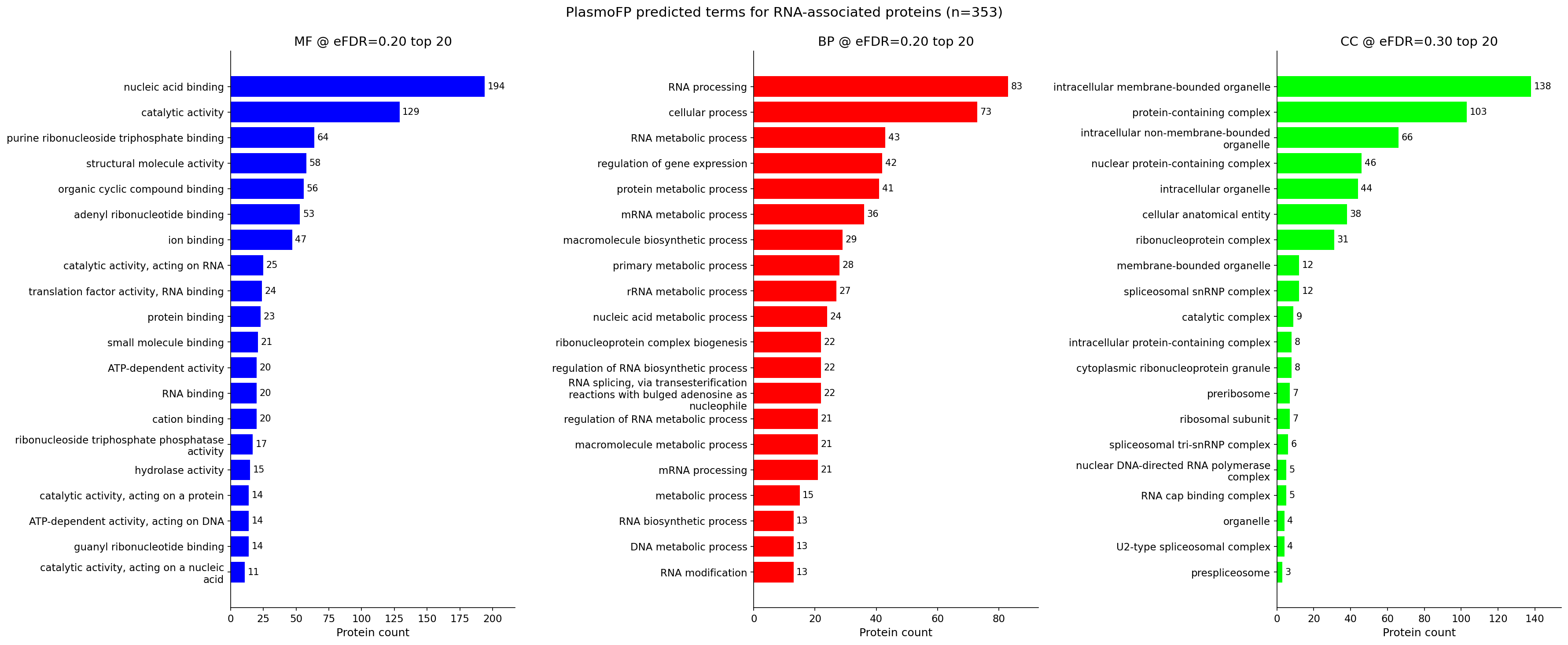

**Figure S9**. **PlasmoFP predicted terms for experimentally validated RNA-associated proteins.** Panels show frequency of predicted terms at specified eFDR thresholds for proteins found in the RNA-associated protein set (n = 353).

A

B

C

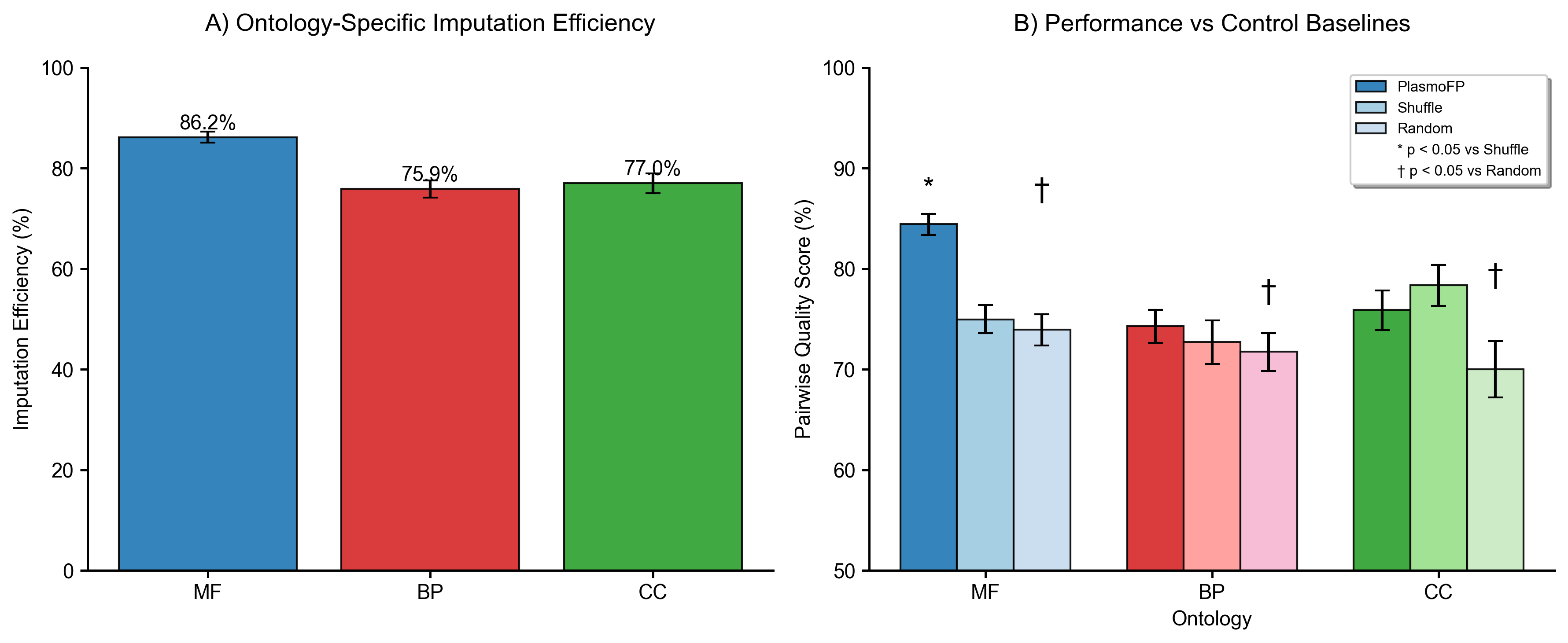

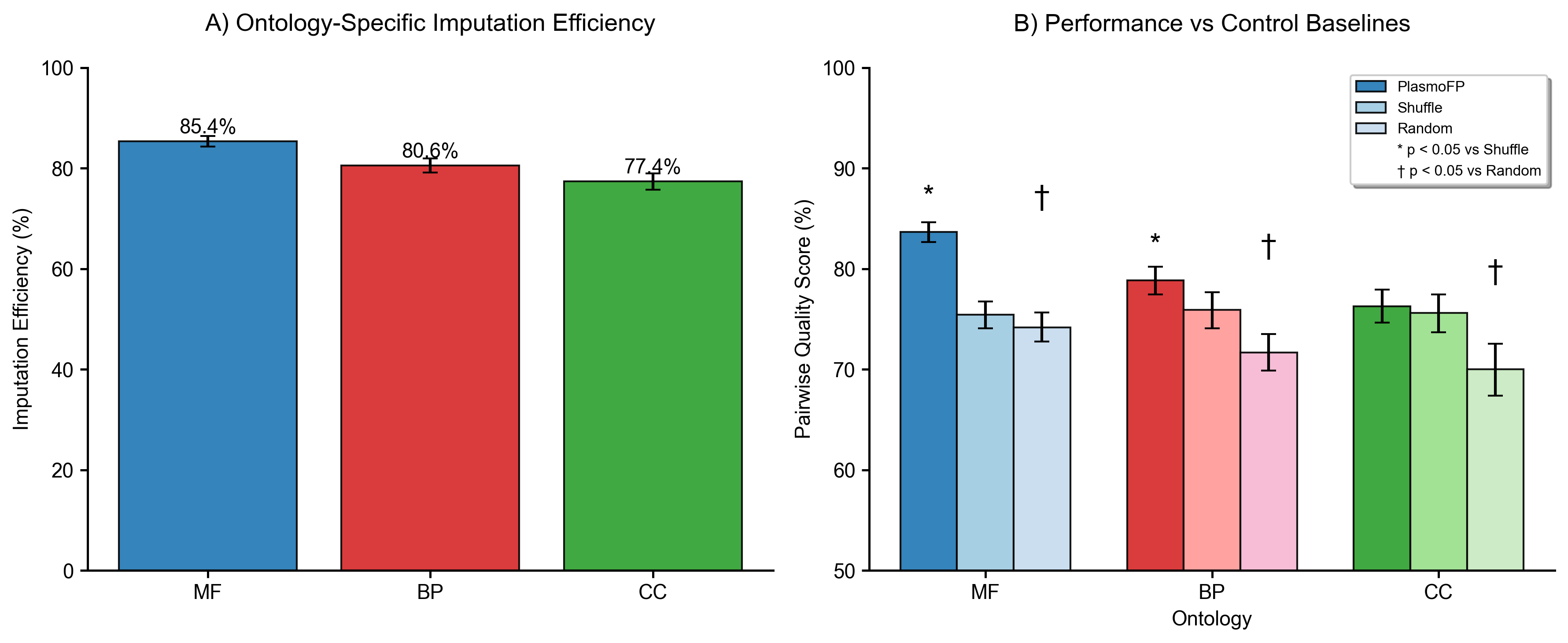

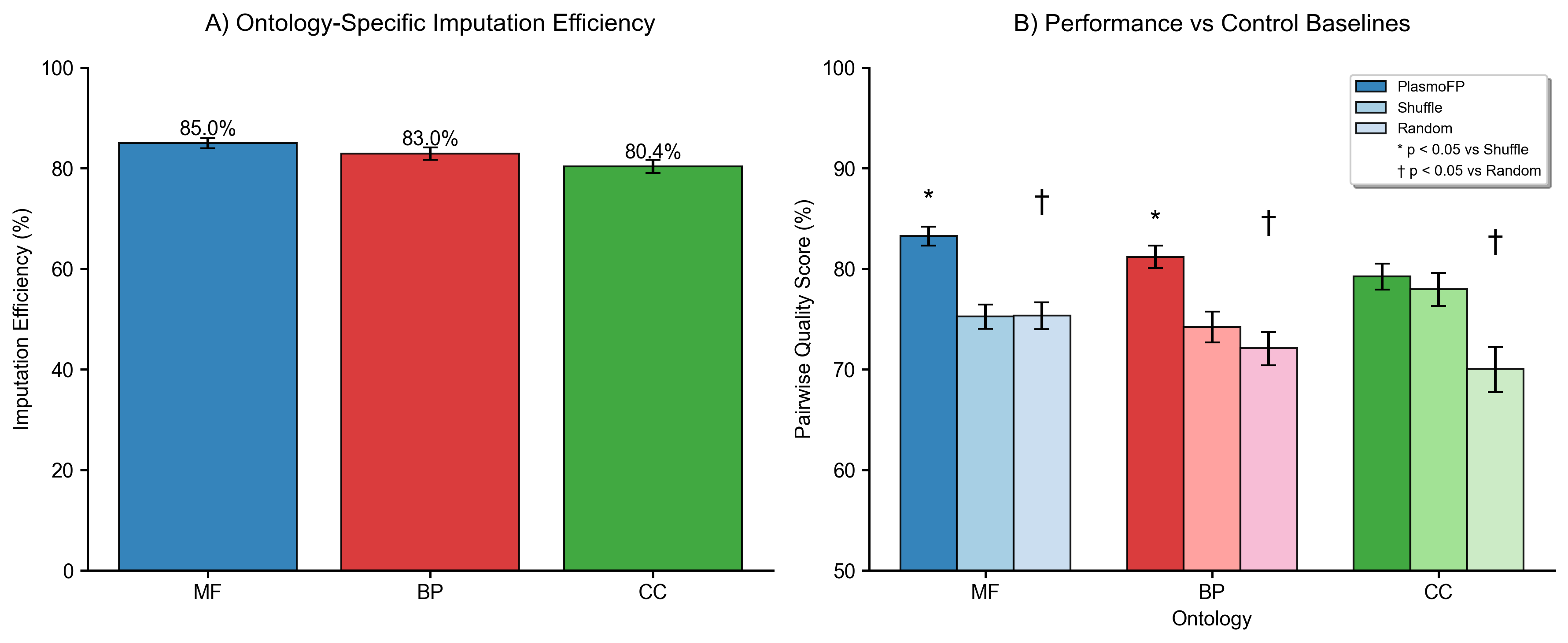

**Figure S10. PlasmoFP performance on an imputation-task for partially annotated proteins.** Cross-ontology imputation efficiency and pairwise quality scores for PlasmoFP at A) 5% eFDR, B) 10% eFDR, and C) 20% eFDR. The right panels show pairwise quality scores for PlasmoFP (dark bars) versus control shuffle baseline (lighter shade) and random (lightest shade). Significance markers: * p < 0.05 vs shuffle; † p < 0.05 vs random (paired permutation over proteins).

A

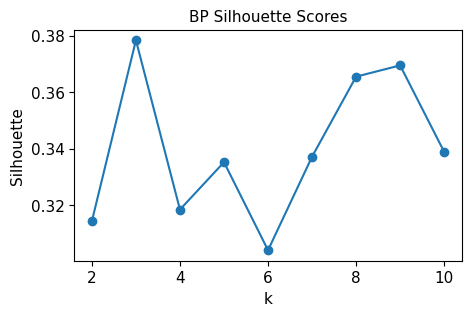

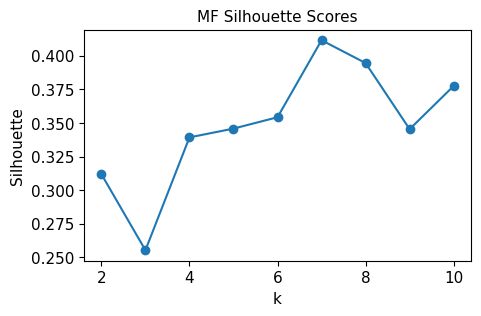

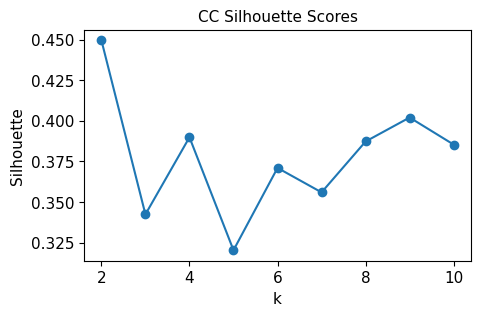

| Ontology  Cluster names | Word clouds of GO term names in each cluster |
| --- | --- |
| **BP**  0: RNA Processing & Modification  1: Cellular & Host Stress Response  2: Regulation of Biological Processes  3: Protein Transport & Localization  4: Amino Acid & Fatty Acid Metabolism  5: Macromolecular Complex Assembly & Organization  6: DNA Damage Response & Repair  7: Nucleotide Metabolism & Biosynthesis | 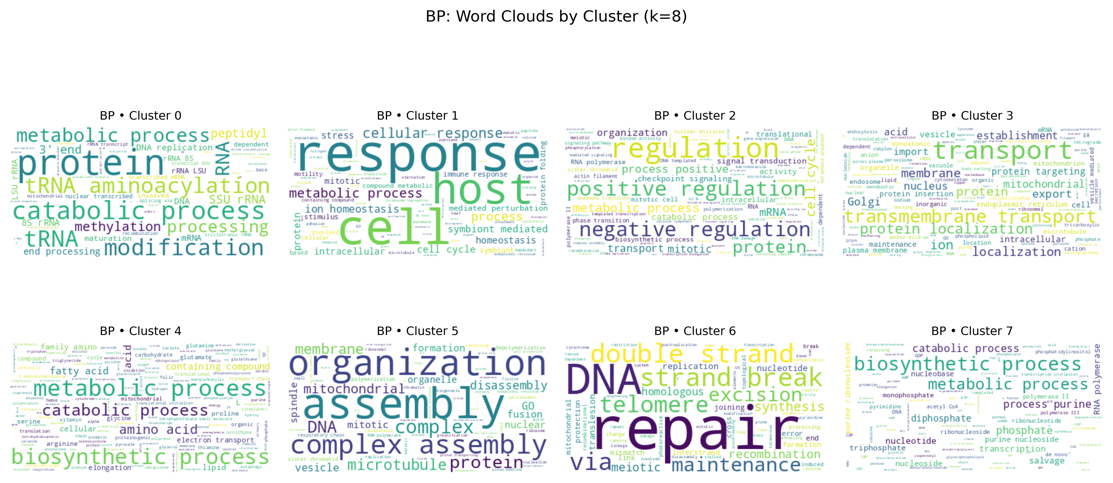 |
| **MF**  0: Redox Activity  1: Hydrolases & Phosphatases  2: DNA/RNA Binding & Polymerase Activity  3: RNA Processing  4: Kinases & Transferases  5: Lyase & Synthase Enzymes  6: Transporter & Channel Activity | 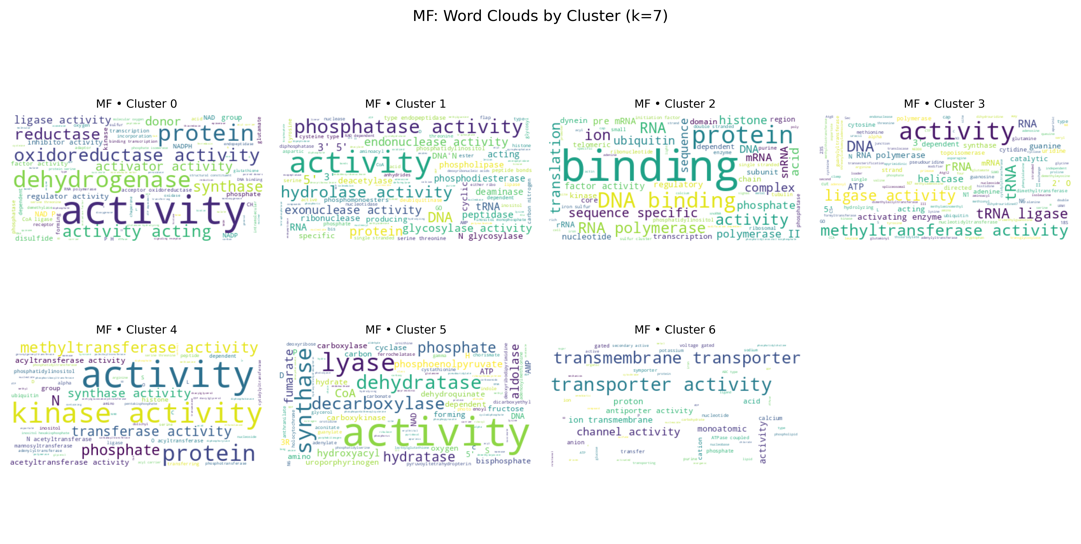 |
| **CC**  0: Membrane & Organelle  1: Gene-Expression Complexes  2: Nuclear Enzyme Complexes  3: Membrane Transport & Vesicle | 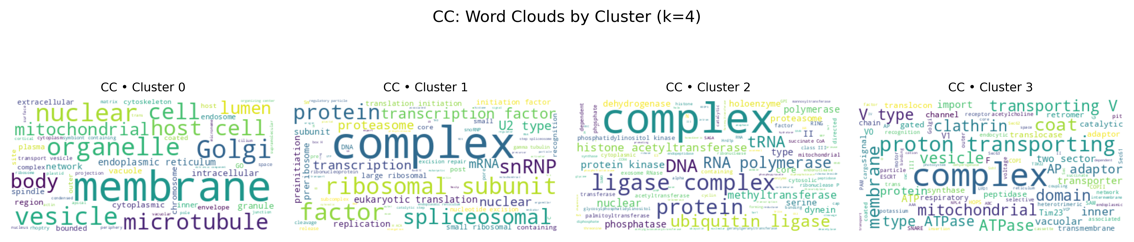 |

B

**Figure S11. GO term clustering.** A) Sillhoute scores of clustering Resnik distance matrix computed on all GO terms found in PlasmoFP predictions and PlasmoDB across all species. B) K-means clusters identified using the Resnik matrix and corresponding cluster names and word clouds.

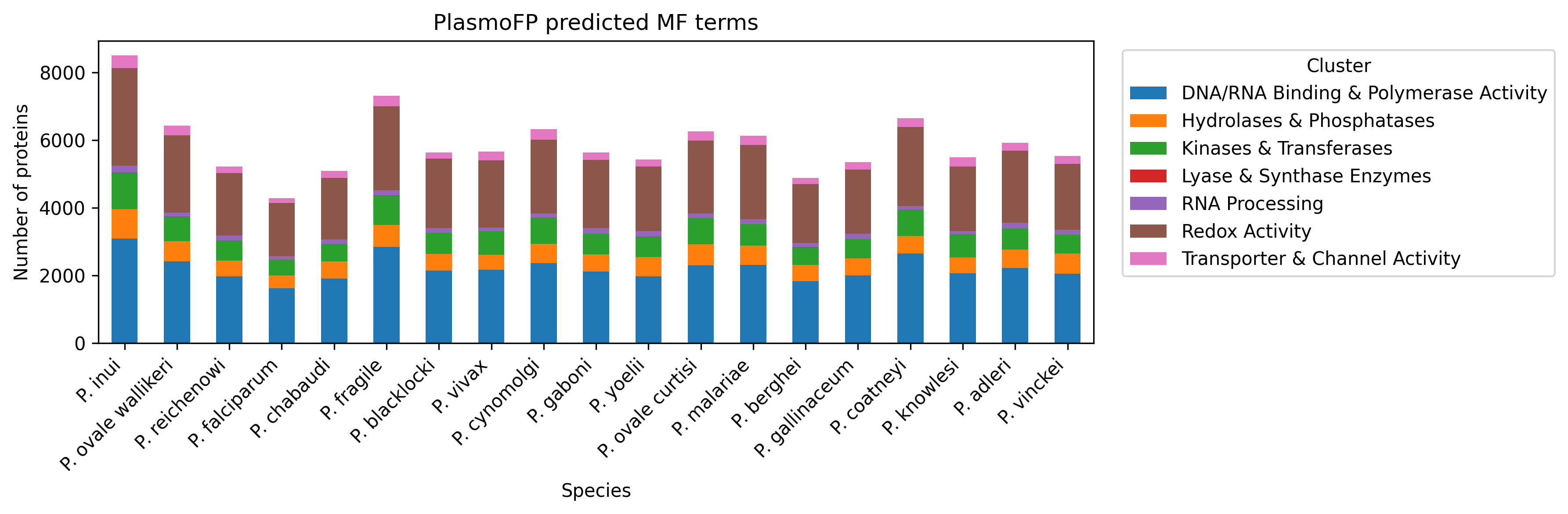

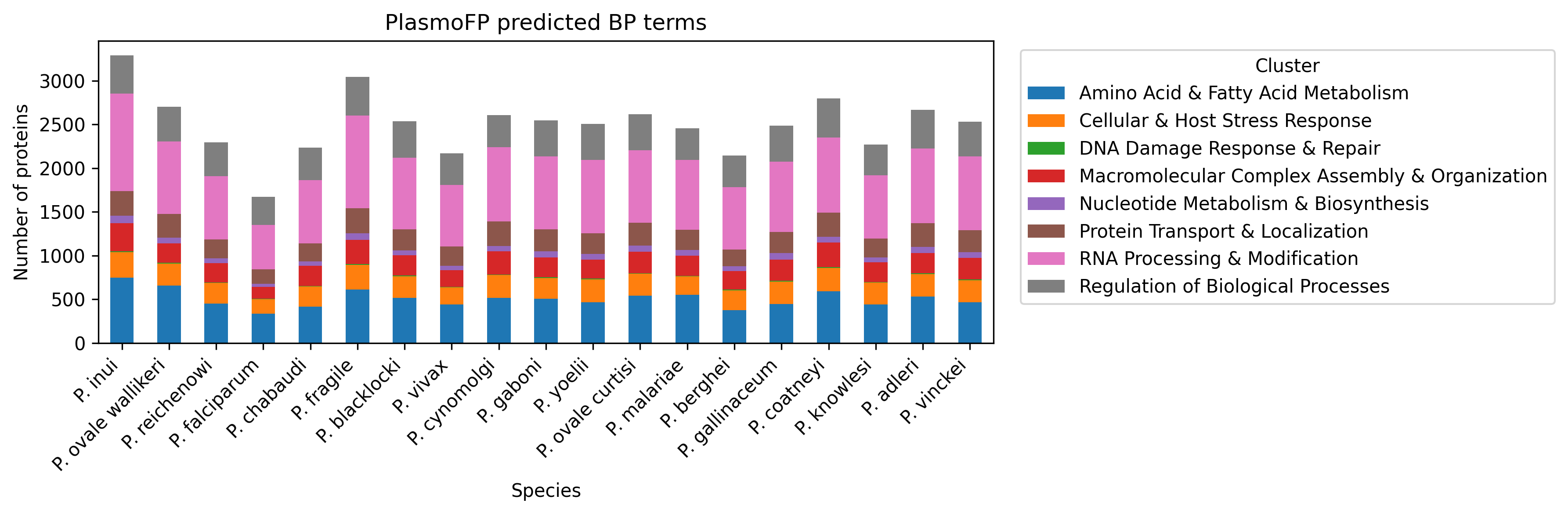

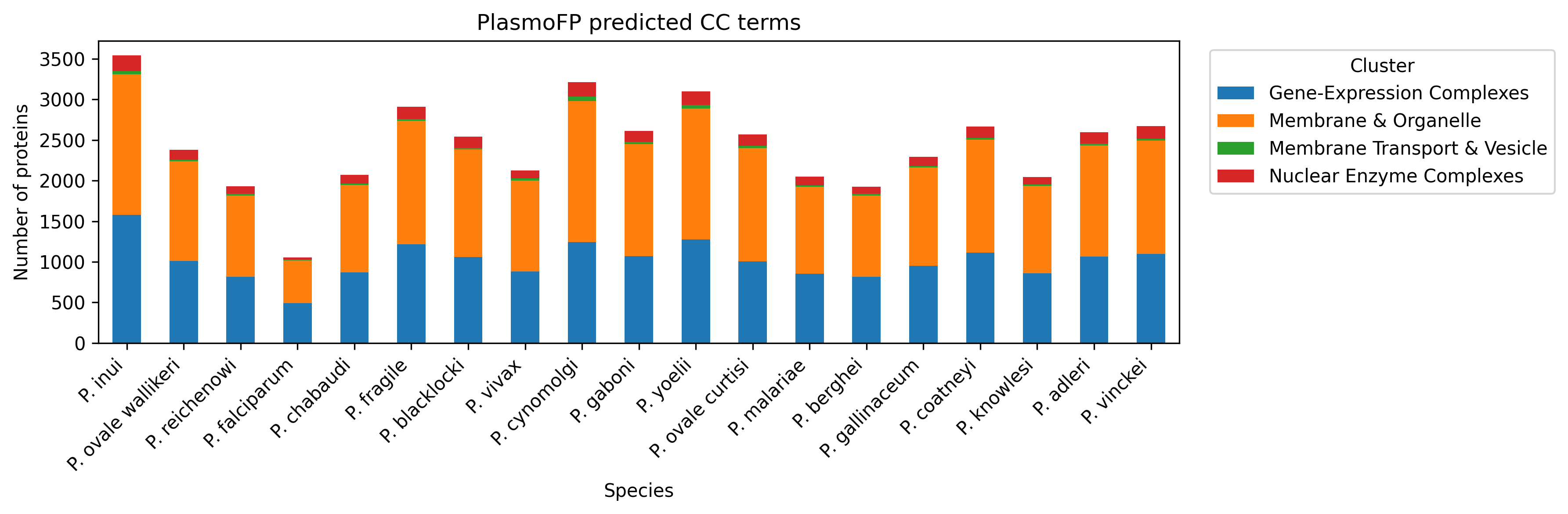

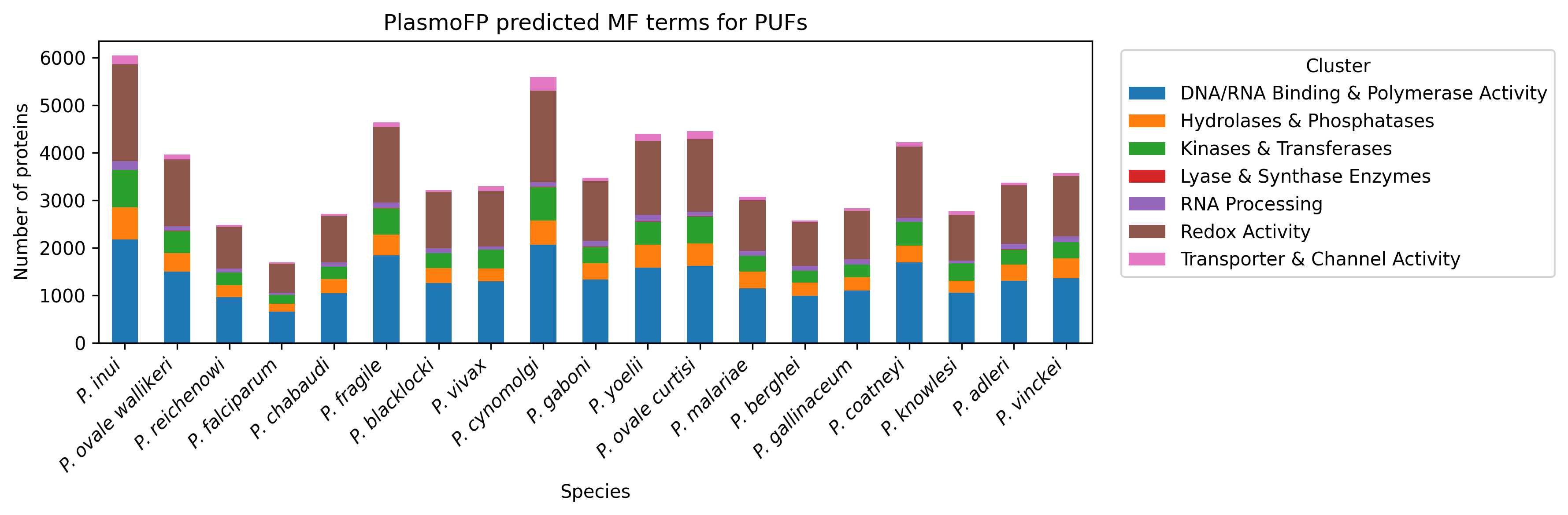

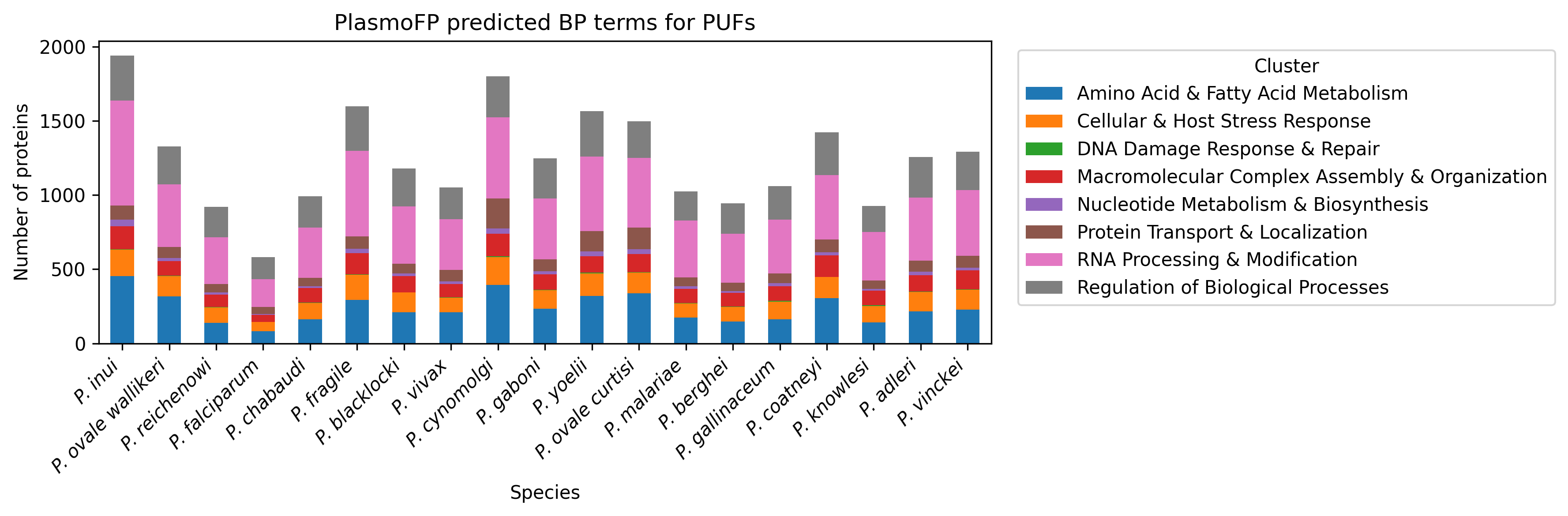

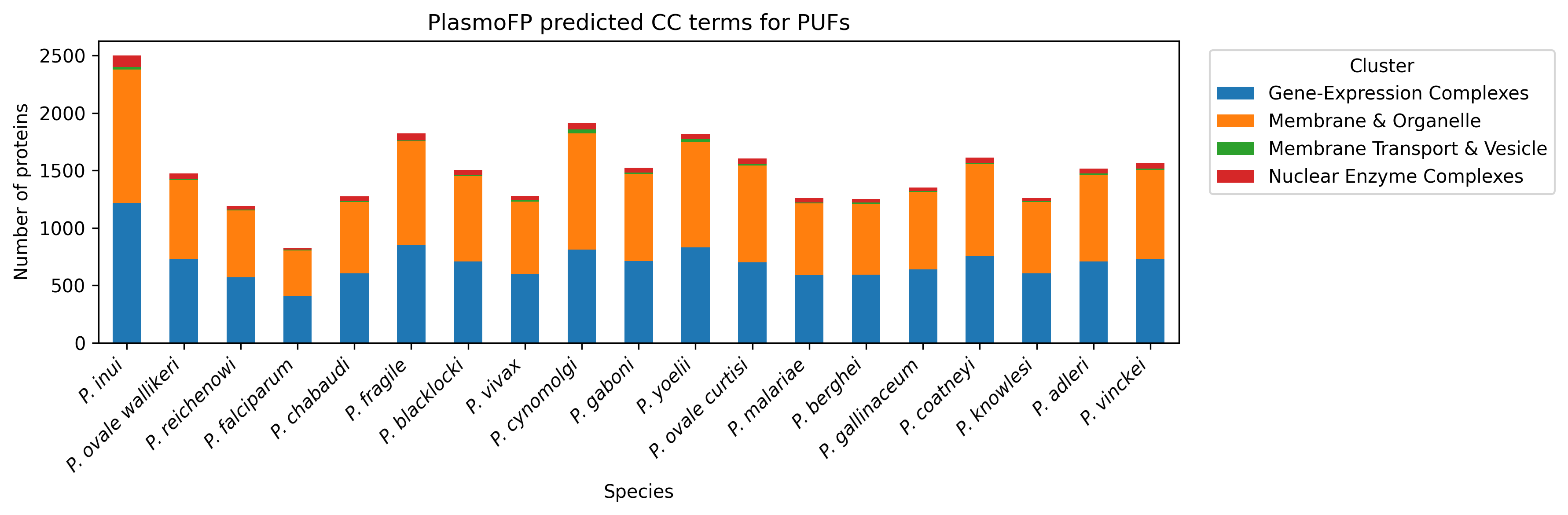

A

B

C

**Figure S12. PlasmoFP predictions for PUFs and partially annotated proteins.** Predictions made by PlasmoFP across all PUFs and partially annotated proteins across A) MF, B) BP, and C) CC subontologies grouped by Resnik defined cluster.

**RNA-associated GO terms:**

GO:0016071: mRNA metabolic process

GO:0006396: RNA processing

GO:0005840: Ribosome

GO:0140098 catalytic activity acting on RNA

GO:0003723: RNA binding

**DNA-associated GO terms**

GO:0003677: DNA binding

GO:0003700: Transcription regulator

GO:0006259: DNA metabolic process

GO:0006260: DNA replication

GO:0006281: DNA repair

GO:0006310: DNA recombination

A

B

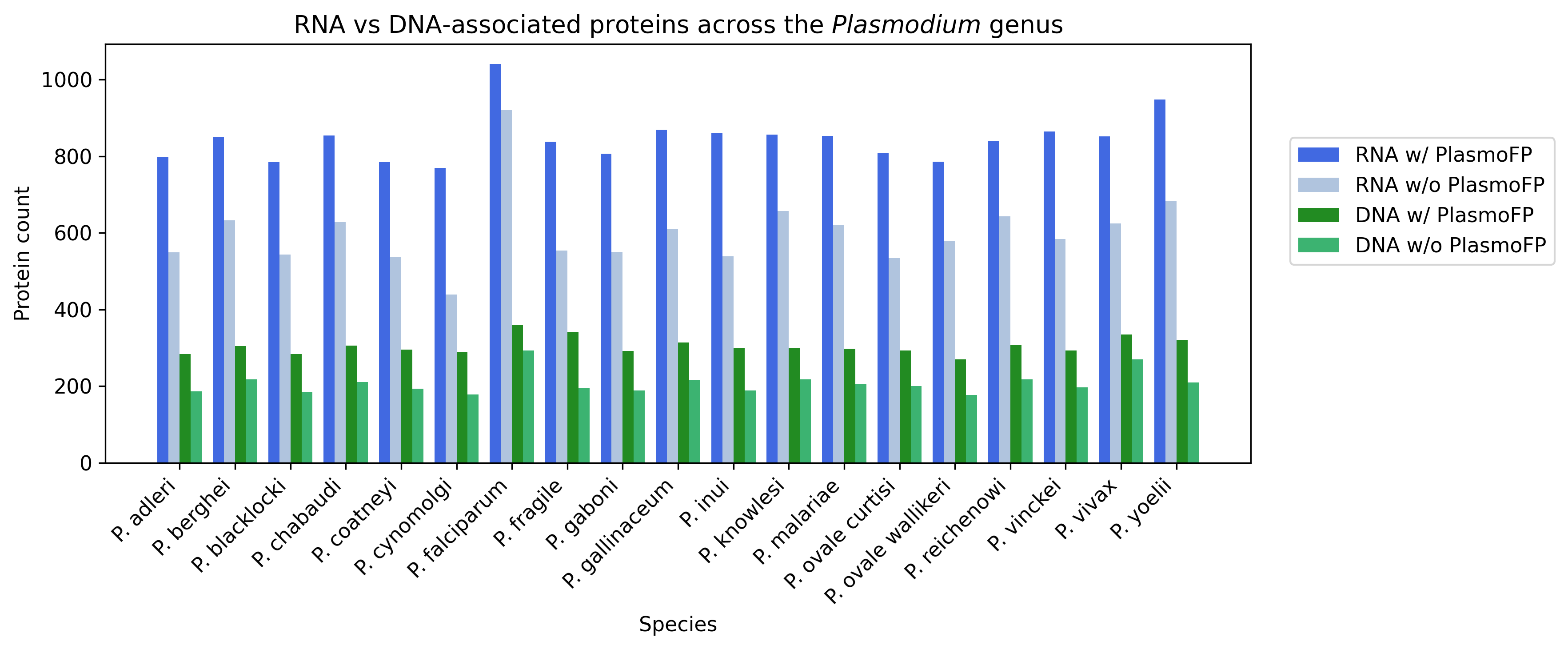

**Figure S13. PlasmoFP expands RNA- and DNA-associated proteins across the *Plasmodium* genus.** A) GO terms used to define “RNA-associated” and “DNA-associated” categories B) For each species, bar heights show the number of proteins annotated as RNA-associated (blue) or DNA-associated (green), before PFP integration (pale bars) and after adding PFP predictions at 30 % eFDR (dark bars).

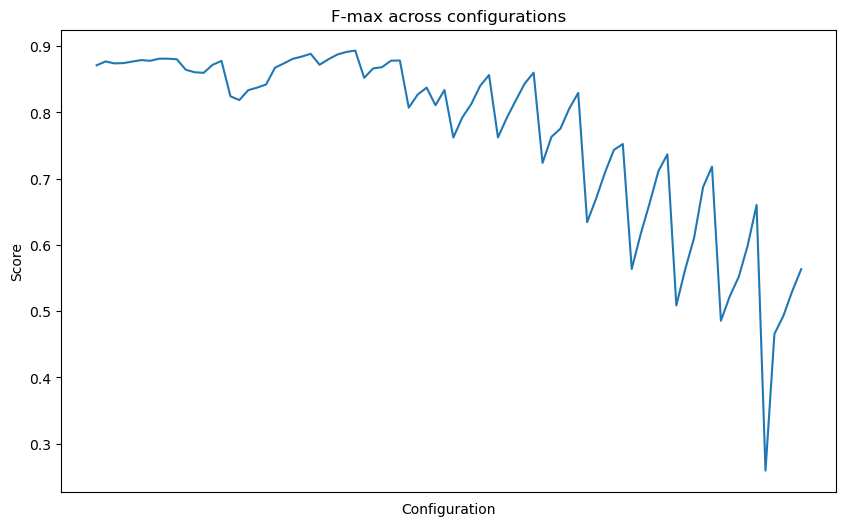

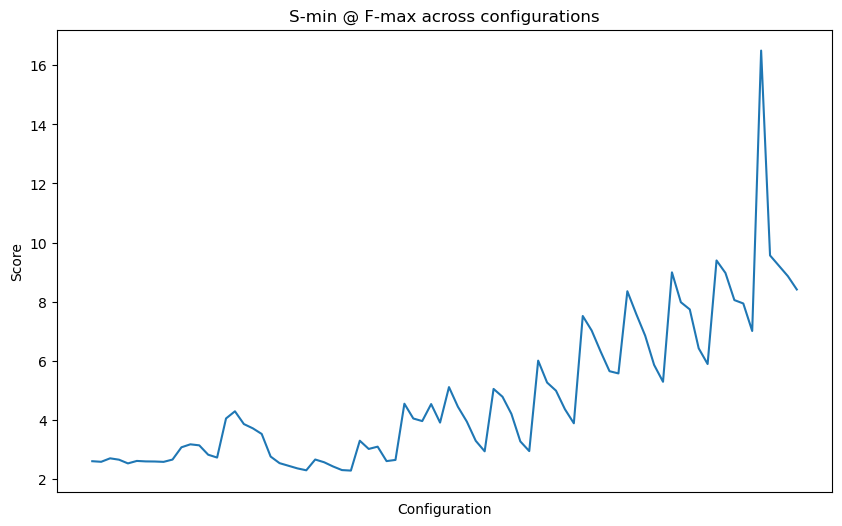

MF

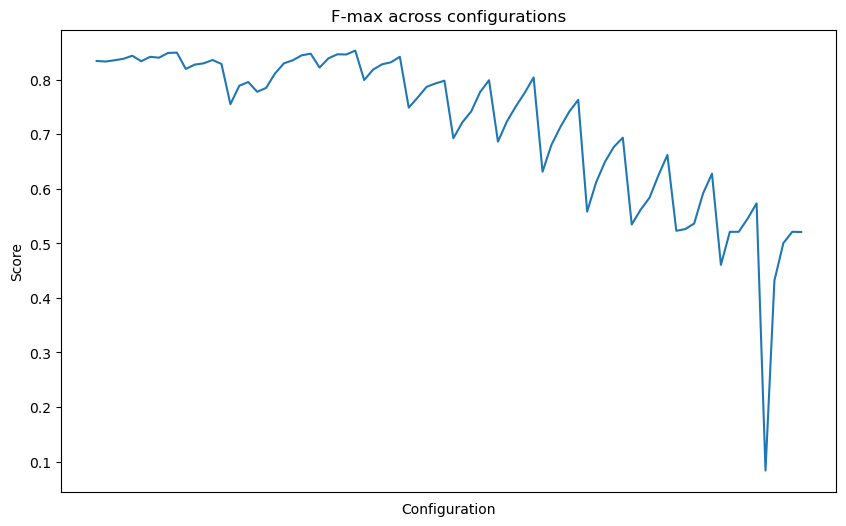

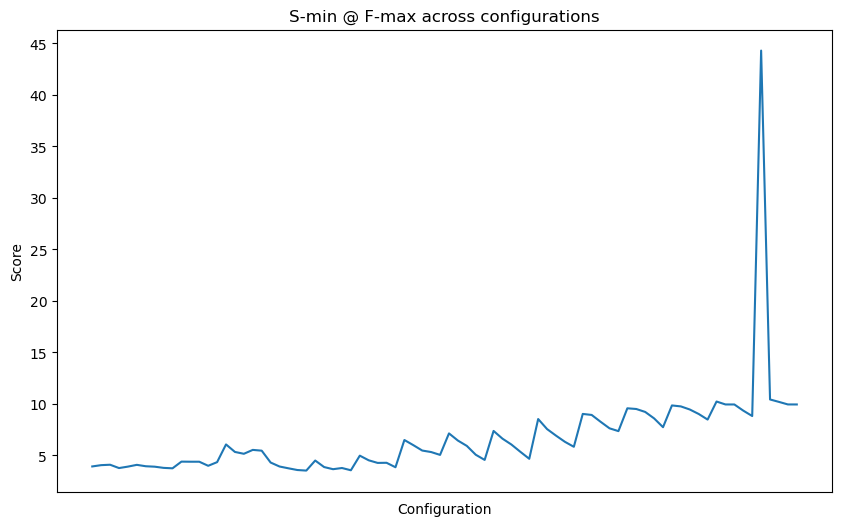

BP

CC

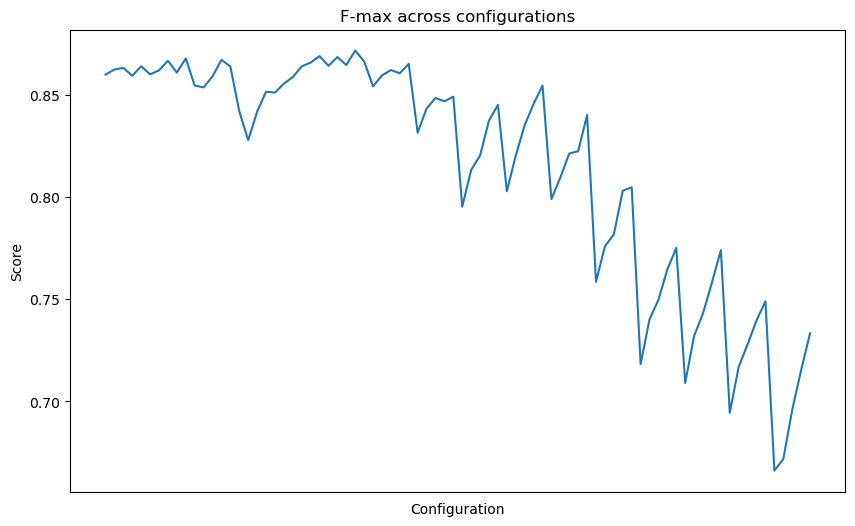

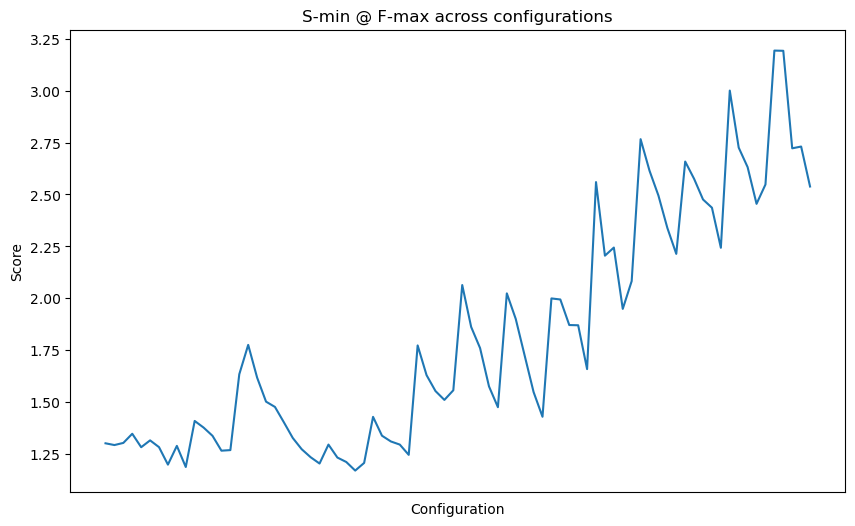

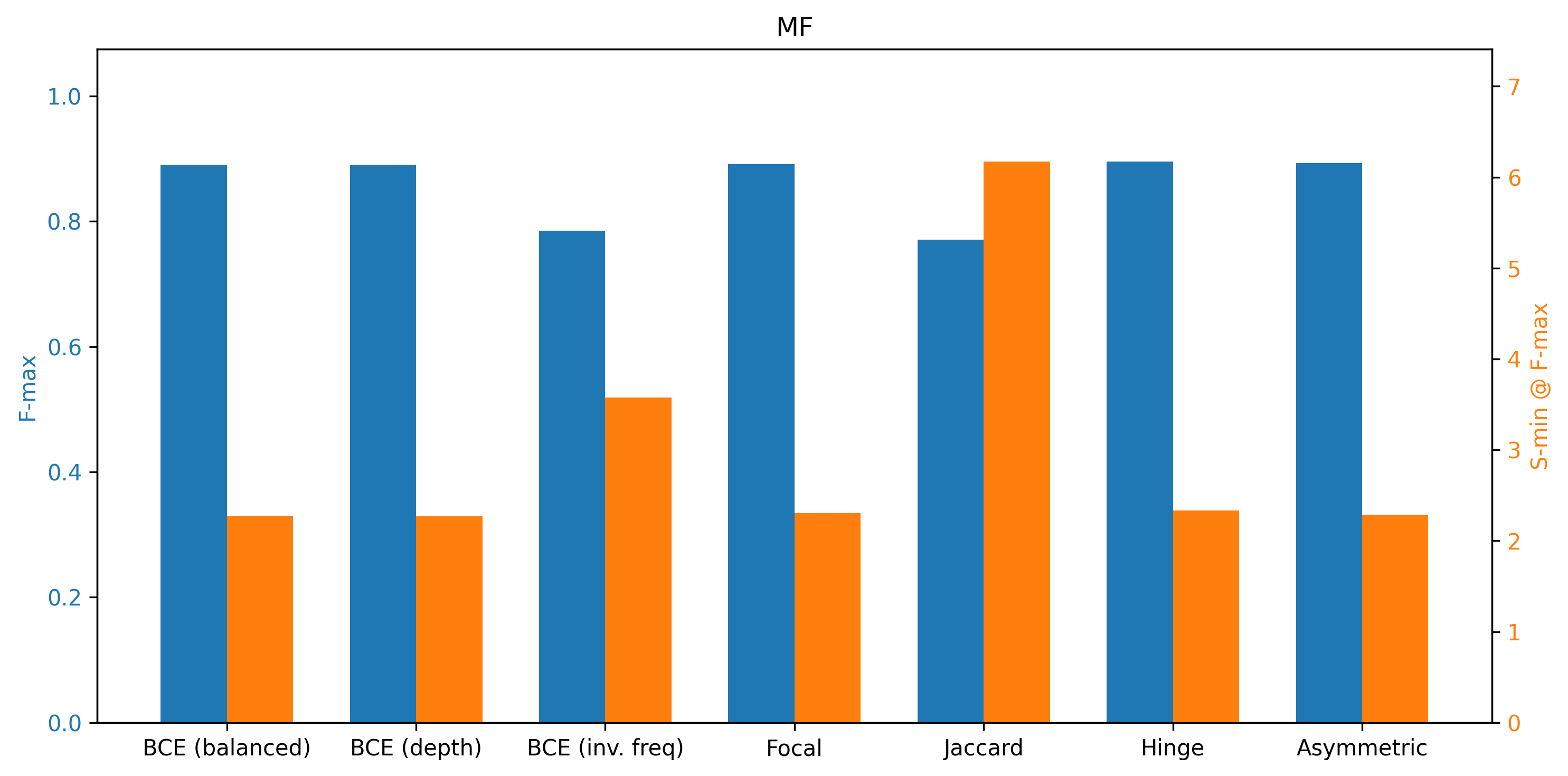

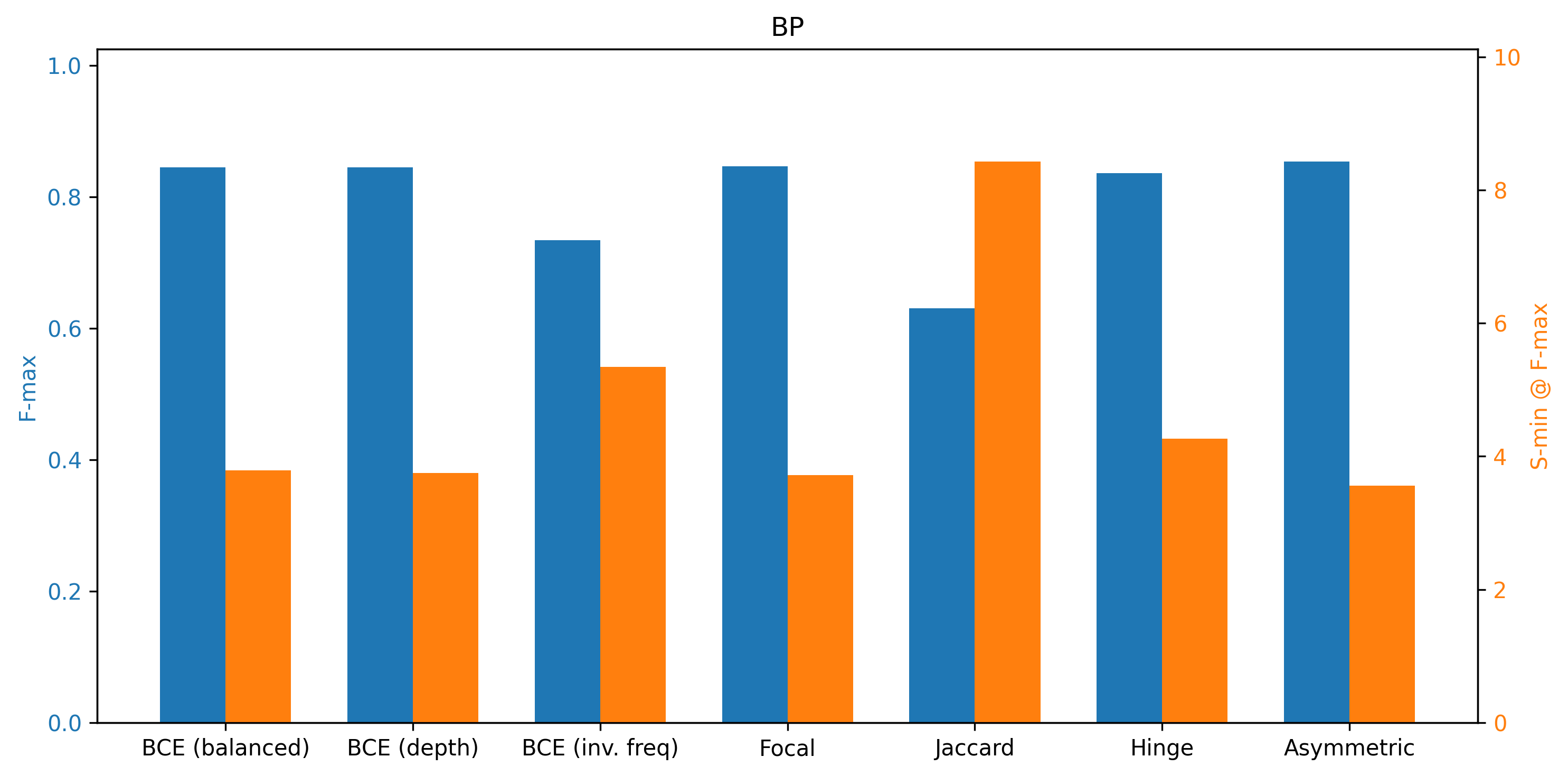

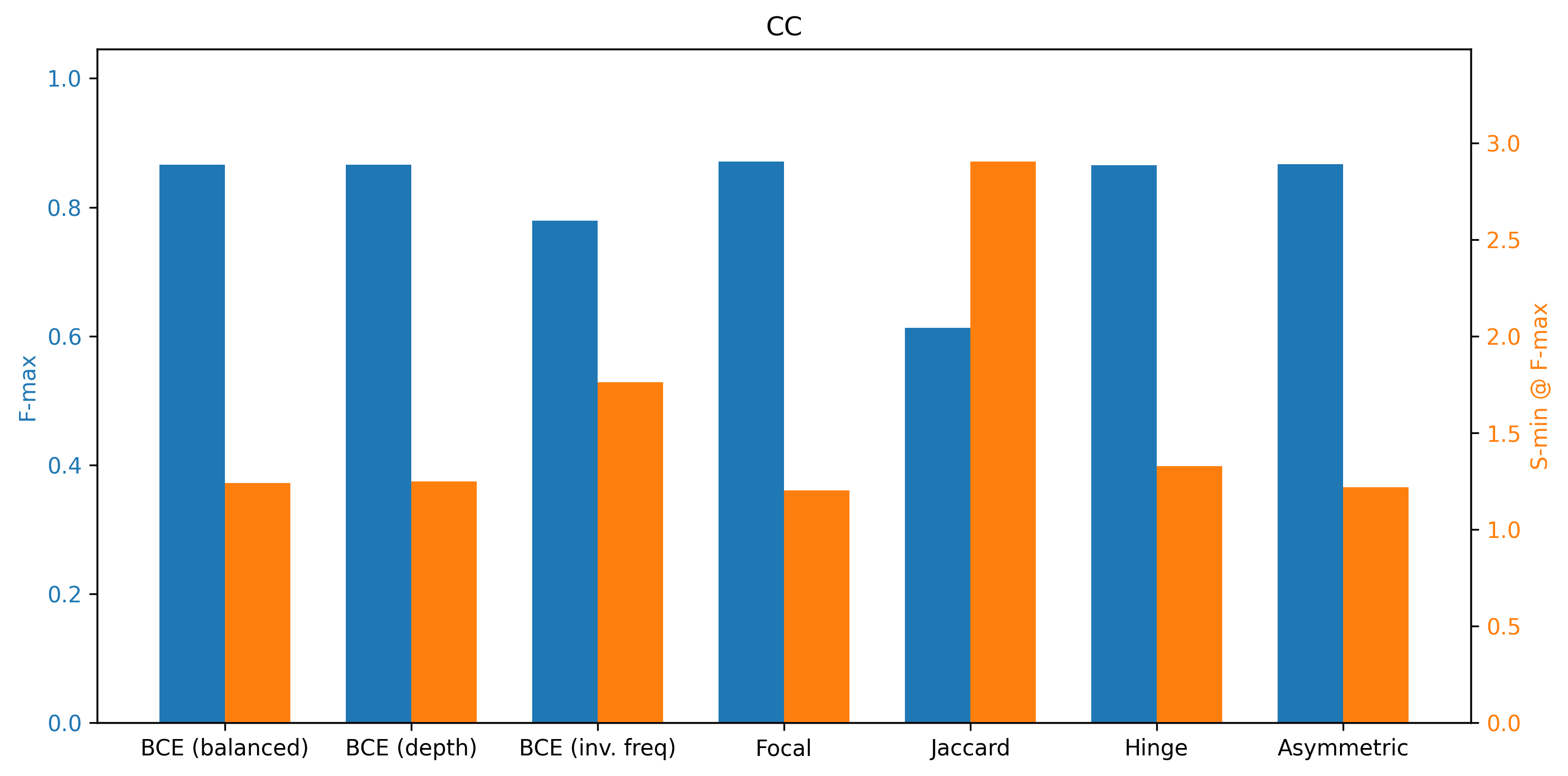

**Figure S14. Hyperparameter tuning.** Tuning results of parameters for PlasmoFP models, organized by ontology. The first column shows model tuning results based on F_max_, the middle column displays results for S_min_, and the final column presents different loss functions applied to the models with the best performance from the first and middle columns.

**Table S1. Nineteen malaria parasite species annotated by PlasmoFP models.** NHP: nonhuman primate.

|  | **Species and strain** | **Host** | **No. predicted protein coding genes** |
| --- | --- | --- | --- |
| 1 | *Plasmodium falciparum* 3D7 | Human | 5318 |
| 2 | *Plasmodium reichenowi* CDC | NHP | 5638 |
| 3 | *Plasmodium adleri* G01 | NHP | 5325 |
| 4 | *Plasmodium blacklocki* G01 | NHP | 5100 |
| 5 | *Plasmodium gaboni* strain G01 | NHP | 5134 |
| 6 | *Plasmodium malariae* UG01 | Human | 5942 |
| 7 | *Plasmodium ovale curtisi* GH01 | Human | 6671 |
| 8 | *Plasmodium ovale wallikeri* PowCR01 | Human | 6228 |
| 9 | *Plasmodium vivax* Salvador 1 | Human | 5530 |
| 10 | *Plasmodium inui* San Antonio 1 | NHP | 5832 |
| 11 | *Plasmodium cynomolgi* M | NHP | 6068 |
| 12 | *Plasmodium knowlesi* strain H | Human, NHP | 5328 |
| 13 | *Plasmodium coatneyi* Hackeri | NHP | 5516 |
| 14 | *Plasmodium fragile* nilgiri | NHP | 5672 |
| 15 | *Plasmodium vinckei brucechwatti* DA | Rodent | 5225 |
| 16 | *Plasmodium yoelii yoelii* 17XNL | Rodent | 5971 |
| 17 | *Plasmodium berghei* ANKA | Rodent | 4945 |
| 18 | *Plasmodium chabaudi chabaudi* AS | Rodent | 5199 |
| 19 | *Plasmodium gallinaceum* 8A | Bird | 5286 |

**Table S2. “Transport” GO term classification.** GO terms found in the BP “Transport” classified into Transporter Classification Database-inspired clusters.

| **GO Term** | **Term Label** | **Cluster** |
| --- | --- | --- |
| GO:0000041 | transition metal ion transport (not explicit mechanism) | Generic/Not Clearly Defined Transmembrane Transport |
| GO:0000054 | ribosomal subunit export from nucleus via nuclear pore | Channel/Pore-based Transmembrane Transport |
| GO:0006405 | RNA export from nucleus via nuclear pore | Channel/Pore-based Transmembrane Transport |
| GO:0006605 | protein targeting (general) | Localization/Targeting without Direct Transmembrane Step |
| GO:0006606 | protein import into nucleus via nuclear pore | Channel/Pore-based Transmembrane Transport |
| GO:0006613 | cotranslational protein targeting to membrane (via translocon channel) | Channel/Pore-based Transmembrane Transport |
| GO:0006811 | monoatomic ion transport usually via ion channels | Channel/Pore-based Transmembrane Transport |
| GO:0006817 | phosphate ion transport often driven by ion gradients | Secondary Active Transmembrane Transport |
| GO:0006839 | mitochondrial transport often via carrier proteins driven by gradients | Secondary Active Transmembrane Transport |
| GO:0006869 | lipid transport may be non-vesicular or carrier-based but unspecified | Generic/Not Clearly Defined Transmembrane Transport |
| GO:0006886 | intracellular protein transport often involves vesicles | Vesicle-mediated Transport |
| GO:0006892 | post-Golgi vesicle-mediated transport | Vesicle-mediated Transport |
| GO:0006913 | nucleocytoplasmic transport via nuclear pore | Channel/Pore-based Transmembrane Transport |
| GO:0008104 | protein localization | Localization/Targeting without Direct Transmembrane Step |
| GO:0008608 | attachment of spindle microtubules to kinetochore is not transmembrane | Localization/Targeting without Direct Transmembrane Step |
| GO:0015031 | protein transport (too broad) | Generic/Not Clearly Defined Transmembrane Transport |
| GO:0015698 | inorganic anion transport often through ion channels | Channel/Pore-based Transmembrane Transport |
| GO:0015711 | organic anion transport often via secondary carriers | Secondary Active Transmembrane Transport |
| GO:0015748 | organophosphate ester transport likely by secondary carriers | Secondary Active Transmembrane Transport |
| GO:0015850 | organic hydroxy compound transport, often via secondary carriers | Secondary Active Transmembrane Transport |
| GO:0015866 | ADP transport (e.g., ADP/ATP exchange in mitochondria) | Secondary Active Transmembrane Transport |
| GO:0015867 | ATP transport (ADP/ATP exchange) | Secondary Active Transmembrane Transport |
| GO:0015988 | energy coupled proton transport against gradient (e.g., ATP-driven) | Primary Active Transmembrane Transport |
| GO:0016192 |  | Vesicle-mediated Transport |
| GO:0016197 | endosomal transport | Vesicle-mediated Transport |
| GO:0016482 | cytosolic transport (movement in cytosol) | Localization/Targeting without Direct Transmembrane Step |
| GO:0031503 | protein-containing complex localization | Localization/Targeting without Direct Transmembrane Step |
| GO:0032509 | endosome transport via MVB | Vesicle-mediated Transport |
| GO:0032940 | secretion by cell (exocytosis) | Vesicle-mediated Transport |
| GO:0033365 | protein localization to organelle (doesn't necessarily imply crossing membrane) | Localization/Targeting without Direct Transmembrane Step |
| GO:0033750 | ribosome localization | Localization/Targeting without Direct Transmembrane Step |
| GO:0034504 | protein localization to nucleus via pore | Channel/Pore-based Transmembrane Transport |
| GO:0035459 | vesicle cargo loading | Vesicle-mediated Transport |
| GO:0035592 | establishment of protein localization to extracellular region (likely secretion pathway) | Localization/Targeting without Direct Transmembrane Step |
| GO:0044743 | protein transmembrane import into intracellular organelle (e.g., via translocon) | Channel/Pore-based Transmembrane Transport |
| GO:0045047 | protein targeting to ER via Sec translocon | Channel/Pore-based Transmembrane Transport |
| GO:0045184 |  | Localization/Targeting without Direct Transmembrane Step |
| GO:0045324 | late endosome to vacuole | Vesicle-mediated Transport |
| GO:0046903 | secretion | Vesicle-mediated Transport |
| GO:0046907 | intracellular transport often involves vesicles | Vesicle-mediated Transport |
| GO:0048193 | Golgi vesicle transport | Vesicle-mediated Transport |
| GO:0050658 | RNA transport (nuclear export) | Channel/Pore-based Transmembrane Transport |
| GO:0051028 | mRNA transport (nuclear pore) | Channel/Pore-based Transmembrane Transport |
| GO:0051168 | nuclear export via pore | Channel/Pore-based Transmembrane Transport |
| GO:0051170 | import into nucleus via pore | Channel/Pore-based Transmembrane Transport |
| GO:0051640 | organelle localization | Localization/Targeting without Direct Transmembrane Step |
| GO:0051649 | establishment of localization in cell | Localization/Targeting without Direct Transmembrane Step |
| GO:0051656 | establishment of organelle localization | Localization/Targeting without Direct Transmembrane Step |
| GO:0051668 | localization within membrane (lateral movement) | Localization/Targeting without Direct Transmembrane Step |
| GO:0055085 | too general | Generic/Not Clearly Defined Transmembrane Transport |
| GO:0070727 | cellular macromolecule localization | Localization/Targeting without Direct Transmembrane Step |
| GO:0070972 | protein localization to ER (not necessarily crossing) | Localization/Targeting without Direct Transmembrane Step |
| GO:0071705 | nitrogen compound transport (unspecified mechanism) | Generic/Not Clearly Defined Transmembrane Transport |
| GO:0071806 | protein transmembrane transport (translocon) | Channel/Pore-based Transmembrane Transport |
| GO:0072530 | purine-containing compound transmembrane transport (likely secondary carriers) | Secondary Active Transmembrane Transport |
| GO:0072594 | establishment of protein localization to organelle | Localization/Targeting without Direct Transmembrane Step |
| GO:0072655 | establishment of protein localization to mitochondrion via import channels | Channel/Pore-based Transmembrane Transport |
| GO:0072657 | protein localization to membrane (not necessarily crossing) | Localization/Targeting without Direct Transmembrane Step |
| GO:0090150 | establishment of protein localization to membrane | Localization/Targeting without Direct Transmembrane Step |
| GO:0098655 | monoatomic cation transmembrane transport (ion channel) | Channel/Pore-based Transmembrane Transport |
| GO:0098657 | import into cell (could be endocytosis or channel) | Generic/Not Clearly Defined Transmembrane Transport |
| GO:0098660 | inorganic ion transmembrane transport | Channel/Pore-based Transmembrane Transport |
| GO:0098661 | inorganic anion transmembrane transport | Channel/Pore-based Transmembrane Transport |
| GO:0098662 | inorganic cation transmembrane transport | Channel/Pore-based Transmembrane Transport |
| GO:0098876 | vesicle-mediated transport to PM | Vesicle-mediated Transport |
| GO:0099515 | actin filament-based transport (not transmembrane) | Cytoskeletal-based Transport |
| GO:0099518 | vesicle cytoskeletal trafficking | Cytoskeletal-based Transport |
| GO:0140056 | organelle localization by membrane tethering | Localization/Targeting without Direct Transmembrane Step |
| GO:0140352 | export from cell (could be secretion) | Generic/Not Clearly Defined Transmembrane Transport |
| GO:0170036 | import into mitochondrion via TOM/TIM (channels) | Channel/Pore-based Transmembrane Transport |
| GO:1901679 | nucleotide transmembrane transport often via carriers | Secondary Active Transmembrane Transport |
| GO:1905039 | carboxylic acid transmembrane transport often secondary | Secondary Active Transmembrane Transport |
| GO:1990542 | mitochondrial transmembrane transport (carrier proteins) | Secondary Active Transmembrane Transport |
